# Exploration of O-GlcNAc-transferase (OGT) glycosylation sites reveals a target sequence compositional bias

**DOI:** 10.1101/2022.09.12.507593

**Authors:** P. Andrew Chong, Michael Nosella, Manasvi Vanama, Roxana Ruiz-Arduengo, Julie D. Forman-Kay

## Abstract

O-GlcNAc transferase (OGT) is an essential glycosylating enzyme that catalyzes the addition of N-acetylglucosamine to serine or threonine residues of nuclear and cytoplasmic proteins. The enzyme glycosylates a broad range of peptide sequences and prediction of glycosylation sites has proven challenging. The lack of an experimentally verified set of polypeptide sequences that are not glycosylated by OGT has made prediction of legitimate glycosylation sites more difficult. Here, we tested a number of intrinsically disordered protein regions as substrates of OGT to establish a set of sequences that are not glycosylated by OGT. The negative data set suggests an amino acid compositional bias for OGT targets. This compositional bias was validated by modifying the amino acid composition of the protein Fused in sarcoma (FUS) to enhance glycosylation. NMR experiments demonstrate that the tetratricopeptide repeat (TPR) region of OGT can bind FUS and that glycosylation-promoting mutations enhance binding. These results provide evidence that the TPR recognizes disordered segments of substrates with particular compositions to promote glycosylation, providing insight into the broad specificity of OGT.

## Introduction

Intracellular *O*-linked-β-*N*-acetylglucosamine (O-GlcNAc) is an essential post-translational modification (PTM). Since discovery of the modification more than three decades ago(1), various proteomic studies have identified thousands of proteins with the modification(2–6). Underscoring the importance of the modification, the single enzyme responsible for adding O-GlcNAc onto proteins, O-GlcNAc transferase (OGT), is required for the viability of dividing mammalian cells and embryogenesis(7, 8). O-GlcNAc is thought to play essential roles in nutrient sensing and stress response with implications for diabetes, cancer and diseases of aging, including neurodegenerative disease(reviewed in (9)). O-GlcNAc-modified proteins implicated in neurodegenerative diseases including amyloid precursor protein(10), tau(11), α-synuclein(12) and superoxide dismutase(13). Mutations in OGT have also been implicated in intellectual disability(14). Unlike other glycan modifications, which are often multimeric(15), O-GlcNAc modification involves addition of a single GlcNAc moiety to serine or threonine hydroxyls. O-GlcNAc significantly affects protein thermodynamic and solvation properties and modulates protein thermal stability(16) and aggregation propensity(11, 17, 18). In addition, O-GlcNAc modification has been recently demonstrated to modulate protein phase separation, based on experimental work with EWS, CAPRIN1 and SynGAP/PSD-95 as well as broader bioinformatics results(19–21).

Given the significant biological impact of this PTM, several studies have grappled with defining OGT sequence specificity(22–26) using O-GlcNAc-modified sites identified in cell extracts by mass spectrometry or by utilizing high throughput assays performed on peptide or protein microarrays(22, 27). O-GlcNAc status *in vivo* is a convolution of multiple factors. These include the specificity of OGT and the specificity of the enzyme that removes O-GlcNAc moieties, O-GlcNAcase (OGA)(28). The efficiency and specificity of OGT is also influenced by the expressed splice isoform(29), since different OGT isoforms contain different numbers of tetratricopeptide (TPR) repeats and TPR repeats are involved in peptide substrate recognition(24, 26, 30–32). OGT may also be recruited to substrates by adaptor proteins(33–35). The concentration of glucose and insulin as well as tissue type and developmental stage also contribute(36, 37). Finally, OGT modification sites are primarily found in extended loops or intrinsically disordered regions (IDRs)(5) that can access the catalytic site.

Several closely related OGT recognition sequences have been identified. For example, Pathak *et al.* identified the OGT recognition sequence as [TS][PT][VT]S/T[RLV][ASY](22). Nevertheless, a scan of O-GlcNAcylated peptide sequences in the PhosphoSite database(38) indicates that most substrates fall outside of this definition, as well as definitions put forward by other groups. Computational methods that make use of machine learning or neural networks to predict sites of O-GlcNAc modification(23, 39–46) have been used to address this shortcoming. These computational methods take both sequence and amino acid combinations into account when making their predicitions. Predictors include YinOYang(43), OGTSite(42), O-GlcNAcscan(46) and O-GlcNAcPRED-II(41), although not all of these predictors are still available online. Evaluating the effectiveness of these predictors is very challenging, in part because of the lack of experimentally verified negative sites. Particular attention must be paid to sensitivity when evaluating these O-GlcNAc predictors, because only a small minority of serines and threonines are expected to be modified. While many proteins can be modified by OGT, the proportion of individual serines and threonines that are modified may be as low as 1.4% (45). The small number of modified residues makes it easy to achieve a high level of accuracy (combined proportion of correctly identified positive and negative sites) by setting a very high threshold for positive site identification. A high threshold enables correct identification of negative sites, which represent the vast majority of sites. The trade-off is that many positive sites will be missed, yielding a low sensitivity (proportion of positive sites correctly identified). Thus, when evaluating O-GlcNAc site predictors, sensitivity is a critical parameter. Taking sensitivity into consideration, O-GlcNAcPRED-II seems to outperform other prediction methods(41, 45). Despite the extensive effort put into these computational methods, they yield many false positive and false negative sites, so experimental validation is still necessary(47, 48).

The inability to clearly define OGT specificity is the result of at least two contributing factors. First, many of these attempts to define specificity have focussed on peptide regions in the immediate vicinity of the glycosylated region, roughly the length of peptide than can be accommodated in the catalytic site of the enzyme. It is now known that efficient substrate recognition can involve a more extended peptide region than what is incorporated into existing predictors(31). For example, efficient glycosylation of the RNA Polymerase C-terminal domain requires more than 20 heptad repeats or over 140 residues(30). The largest isoform of OGT (ncOGT) is comprised of a catalytic domain preceded in sequence by 13 tetratricopeptide (TPR) repeats (24, 49, 50). The TPR region forms a large superhelix that has been shown to influence recognition of extended lengths of substrate peptides(30, 51, 52). Thus, improving predictors will require consideration of a broader sequence context.

Secondly, the inability to clearly define OGT specificity is due to the lack of an experimentally verified negative data set for prediction purposes. Existing predictors have used protein regions not annotated as glycosylated, but found in proteins that are glycosylated by OGT, as a negative dataset(39, 42), or alternatively human proteins from the UniProt database(53) that are not explicitly known to be glycosylated or predicted to be glycosylated(40). Taking protein regions not annotated as glycosylated, but found in proteins that are glycosylated has the advantage that the proteins are known to exist spatially and temporally near to OGT, an important consideration(54). Nevertheless, the assumption that the absence of mass spectrometry data supporting glycosylation at a specific site is evidence that the site is not glycosylated is a poor one. Mass spectrometry studies often do not achieve full coverage for a particular protein and certainly not for the entire proteome(55). Coverage of the proteome in higher organisms typically does not exceed 10%. In one study where over 1500 O-GlcNAc modified proteins were identified, modification sites could only be assigned in 80 proteins(3). Even when a high degree of coverage is achieved, the identification of post-translational modification sites depends heavily on the abundance of a particular protein(5), especially when the sites are sub-stoichiometrically modified as is often the case for O-GlcNAc. Finally, identification of specific sites is hindered by the lability of the O-GlcNAc modification in MS/MS experiments(56, 57). For these reasons, construction of a negative dataset based on sites that are not annotated as glycosylated is not a good strategy. In one recent study the majority of proteins identified as being O-GlcNAc modified were not previously known to be modified(16). The case of Lamin A serves to illustrate the point. Prior to 2018, Lamin A was shown to have two glycosylated sites(57); more recently, an additional nine sites were identified(47). Only two of the Lamin A sites (S612 and T643) are currently listed in the PhosphoSite O-GlcNAc database. Interestingly, the double mutant S612A/T643A of Lamin A is as robustly O-GlcNAc modified as the WT protein. Inclusion of the nine additional sites in the negative database would clearly impair prediction. The Ewing sarcoma protein (EWS) is another excellent illustration, since it is a well-documented OGT substrate(58) but is not present in the PhosphoSite O-GlcNAc database which was used as the basis for most of the existing predictors.

Here we explored the utility of an experimentally verified negative dataset for prediction of likely glycosylation target sites for the longest isoform of OGT. Initial work on glycosylation of the three FET family proteins, Fused in sarcoma (FUS), EWS and TATA binding protein-associated factor 15 (TAF15), provided insight and suggested a way forward for site prediction. As in previous examples, we used the PhosphoSite O-GlcNAc database as our positive set, but extended the length of the peptide region considered. To obtain a negative dataset, we purified a set of IDRs and subjected them to optimized glycosylation reactions with purified recombinant OGT, then used whole protein mass spectrometry rather than MS/MS to identify proteins that were not being glycosylated. We computationally optimized a scoring matrix to distinguish between the positive set and our experimental negative dataset. The scoring matrix suggests that OGT substrates have an amino acid compositional bias that extends beyond the polypeptide region that can be accommodated in the catalytic site of the enzyme. Compositional mutants of the FUS N-terminal low-complexity region (LCRN) support an OGT compositional bias. We verified that the TPR region of OGT (OGT-TPR) can bind to FUS and an enhanced-glycosylation mutant of FUS, leading us to speculate that the TPR region enables OGT to glycosylate substrates that are not optimally recognized by the catalytic site.

## Results

### Glycosylation Stoichiometry of the FET Proteins

Previously, Kamemura reported that, of the three FET proteins FUS, EWS and TAF15, only EWS is glycosylated with high stoichiometry by OGT(58). Since the FET proteins are homologous, this suggested that further analysis of FET protein glycosylation might inform our understanding of OGT specificity. We assayed glycosylation levels on undigested protein samples using electrospray ionization mass spectrometry following *in vitro* glycosylation of purified human FUS, EWS and TAF15 fragments (Fig. 1a-c). Specifically, we measured glycosylation of the N-terminal low-complexity (LCR_N_) regions of these FET proteins, since this is the region of EWS which is glycosylated(19, 59): FUS (aa 1-214), EWS (aa 1-264), TAF15 (aa 1-210), hereafter referred to as FUS, EWS and TAF15, respectively. A distribution of glycosylated states with three to ten added O-GlcNAc moieties was observed for EWS. No unglycosylated EWS was observed following the reaction. In contrast, the majority of the TAF15 LCRN protein was not glycosylated, although a small amount of singly glycosylated protein was observed. Modest glycosylation of FUS LCRN was observed, with a median of two sugars groups added. These experiments confirm that EWS can be heavily glycosylated, while FUS is modestly glycosylated and TAF15 is largely unglycosylated *in vitro.*

**Figure 1.**
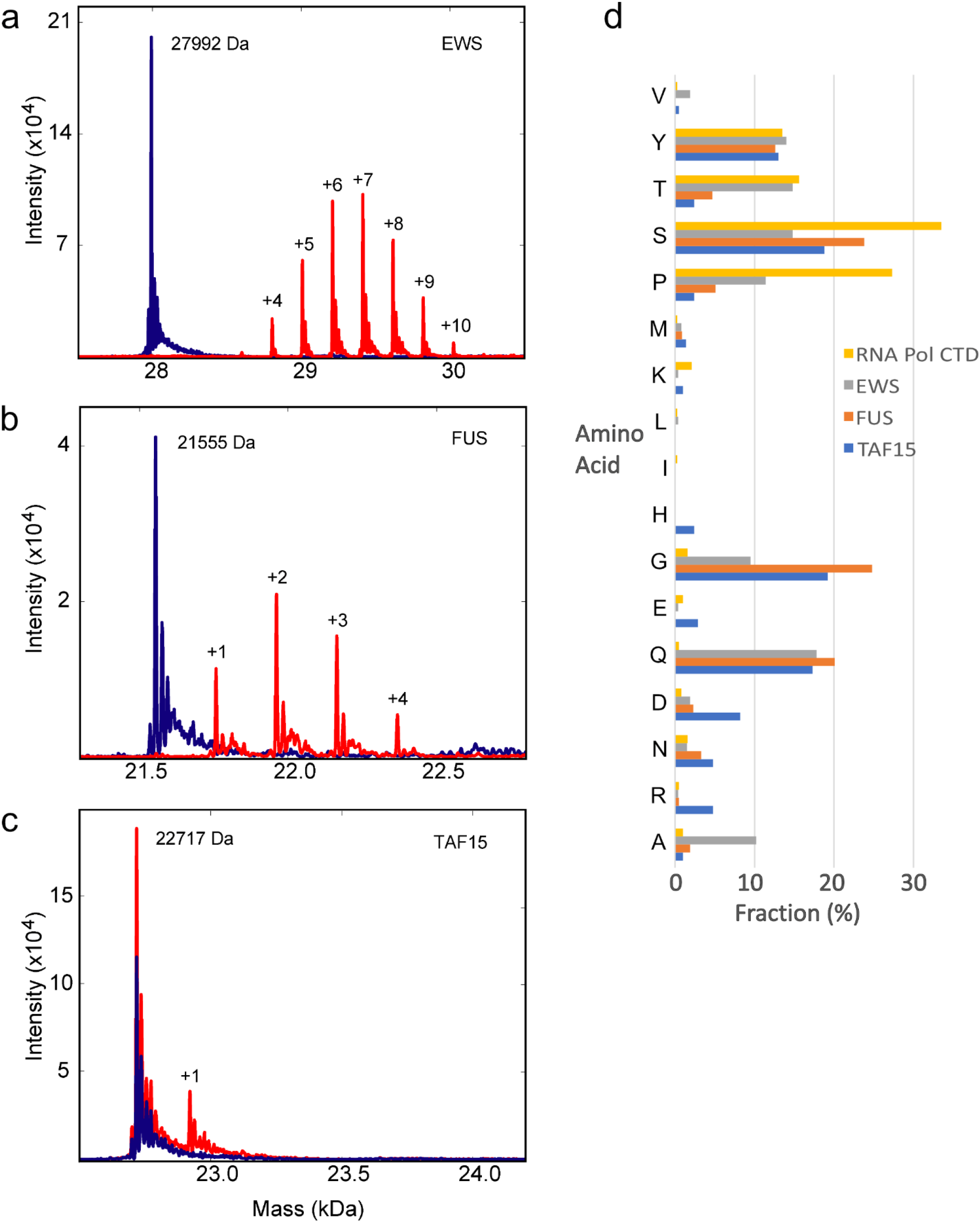
FET protein glycosylation by OGT. Intact mass spectrometry profiles of the LCR_N_ regions of (a) EWS, (b) FUS and (c) TAF15 before and after glycosylation by OGT. Spectra of the unglycosylated proteins are shown in blue and spectra following glycosylation (5 hours at lab temperature) are shown in red, with the number of added sugar groups shown. (d) Amino acid composition of FET protein LCR_N_ and RNA Polymerase II CTD. Compositions are shown for FUS (aa 1-214), TAF15 (aa 1-210) and EWS (aa 1-264) and RNA Polymerase II subunit RPB1 (aa 1586-1970) as labelled in the legend. Only amino acids that are observed in at least one of these regions are shown.

### Glycosylation Sites in EWS

To identify specific EWS glycosylation sites, we subjected the EWS LCR_N_ to chymotrypsin digestion followed by LC-MS/MS. As observed by others, we found that the O-GlcNAc group is labile and is removed during the peptide fragmentation step. Thus, we could determine which peptides are glycosylated, but could not identify specific serines and threonines that are glycosylated. Glycosylation sites were spread across the LCRN as indicated in Table 1. We can determine that there are more than 14 glycosylation sites, though we never observed EWS protein modified at 14 or more sites simultaneously. We speculate that glycosylation at some sites might inhibit glycosylation at nearby sites, thus making it unlikely to observe EWS glycosylated at all possible sites.

**Table 1.** Location of Glycosylation Sites in the EWS LCR_N_. Peptides identified by the MS-MS sequence are listed. The number of glycosylation sites for each peptide was determined by the difference in the molecular weight of the parent ion. LC-MS/MS data: DOI: 10.5281/zenodo.6986306.

| EWS peptide span | Observed glycosylation sites (total number of serines and threonines in peptide) |
| --- | --- |
| 45-61 | 3 (6) |
| 87-112 | 3 (10) |
| 119-158 | 2 (9) |
| 145-170 | 2 (8) |
| 171-195 | 2 (6) |
| 209-226 | 1 (6) |
| 230-254 | 3 (8) |
| 248-264 | 1 (6) |

### Amino Acid Variation in the FET LC Regions

The stark difference in FET protein LCR_N_ glycosylation stoichiometry was surprising since their sequences share many features. These LCR_N_ are primarily comprised of the amino acids glycine, alanine, serine, threonine, tyrosine, proline and glutamine, a composition that is similar to the RNA polymerase C-terminal domain (RNA Pol CTD), which is also known to be glycosylated by OGT. However detailed comparison of the sequences suggests explanations for their differing substrate specificity (Fig. 1d). EWS has a much higher percentage of alanine, proline and threonine residues than either FUS or TAF15. The fractional content of glycine and serine is lower for EWS than either FUS or TAF15. The RNA Pol CTD is noticeably depleted in glycine and glutamine residues relative to the FET proteins. TAF15 has a notably higher proportion of charged residues including arginine, aspartic acid and glutamic acid. Observing these differences, we decided to investigate the importance of amino acid composition outside of the immediate vicinity of the glycosylated sites.

### Computational Optimization of OGT Substrate Prediction

The inability of OGT to appreciably glycosylate TAF15 suggested that it would be possible to develop an experimentally verified negative dataset to improve substrate prediction. To that end, a number of known IDRs were subjected to glycosylation under optimal conditions followed by intact mass spectrometry. The EWS LCR_N_ served as a positive control for this experiment. High quality mass spectra for IDRs of SARA (aa 766-822, human)(60), DDX4 (aa 1-236, mouse)(61), TAF15 (aa 1-210, human), CFTR (aa 654-838, human)(62) and FMRP (aa 445-632, human)(63) demonstrated that these proteins were not glycosylated to any appreciable extent even after 15 hours of reaction time(Fig. 2). Only one of the IDRs that we tested, from the yeast protein Sic1 (aa 1-90)(64), was significantly glycosylated under our reaction conditions (not shown) and thus excluded from our negative data set. (Hereafter, these IDRs are referred to only by the name of the protein.) Close inspection of the DDX4 and TAF15 spectra showed a small fraction of protein glycosylation on a single site. Nevertheless, we decided to keep DDX4 and TAF15 in our negative data set, since the site(s) are clearly less than optimal and barely glycosylated even following overnight incubation. All peptides centered on serine or threonine residues were extracted from the relevant TAF15, CFTR, DDX4, FMRP and SARA sequences to use as a negative dataset. The negative set includes a total of 135 peptides, though many contain overlapping sequences. Because of the small size of the dataset, we did not exclude any of the peptides for subsequent testing of the approach. Our positive dataset consists of the sites listed in the PhosphoSite O-GlcNAc database, which includes 1830 sites primarily from mouse, human and rat proteins.

**Figure 2.**
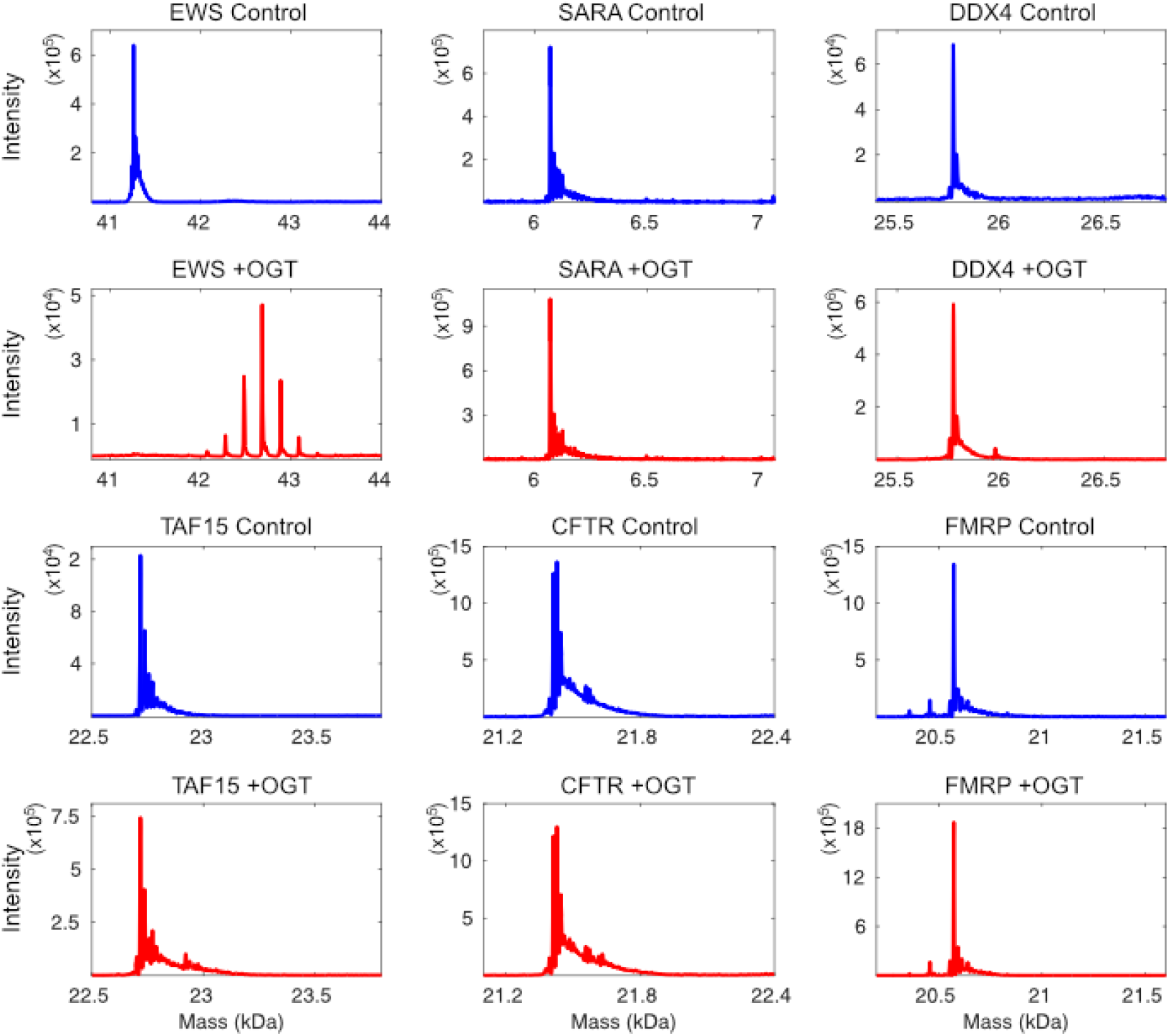
Development of an experimental negative dataset for OGT glycosylation. Mass spectra of EWS (positive control), SARA (aa 766-822), DDX4 (aa 1-236), TAF15 (aa 1-210), CFTR (aa 654-838) and FMRP (aa 445-632) before (blue) and after (red) overnight glycosylation with OGT.

A simple computational strategy was used to optimize a substrate scoring matrix (Fig. 3a,b). The substrate scoring matrix had position relative to the potential glycosylation site along one axis and the amino acids along the other axis. Tryptophan and cysteine were excluded from the scoring matrix, as they were considered too rare to properly evaluate. Substrates in the positive and negative sets were scored by summing the value of the appropriate matrix positions for each amino acid in the sequence. Random modifications were made to the matrix and were kept if the distance between the median scores of the positive and negative dataset increased. The optimized matrices converged on similar matrices, despite starting from very different starting matrices. Matrices for peptide lengths of 15, 23, 31 and 39 were evaluated. As some of the observed trends spanned the longest 39 residue peptides, we chose this length for our further work, consistent with longer lengths being required for optimal glycosylation of some substrates.

**Figure 3.**
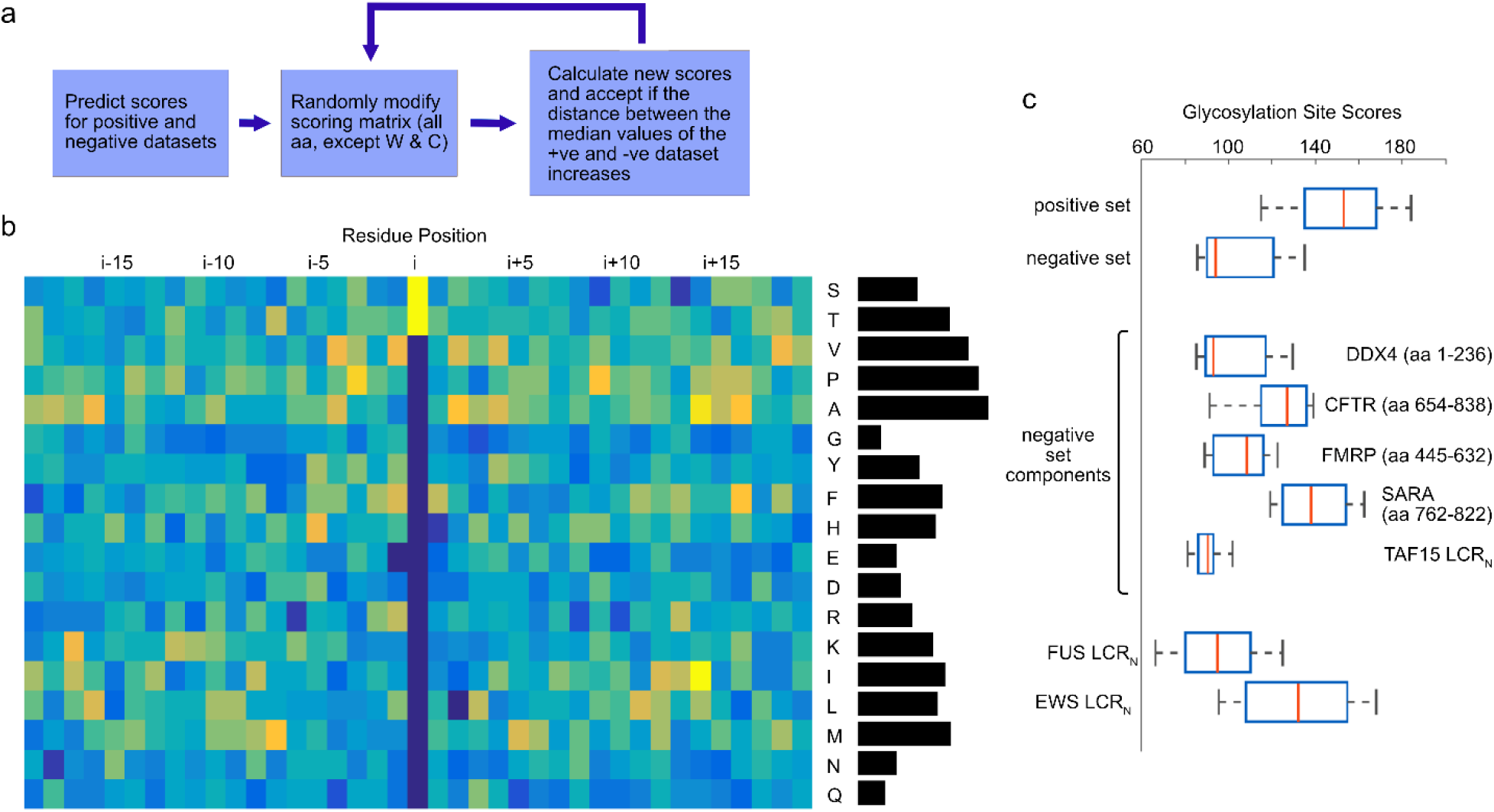
Optimization of a matrix for prediction of OGT glycosylation sites. (a) Schematic for the optimization process. (b) Optimized matrix, with residue position along the horizontal axis and amino acid type along the vertical axis. Yellow represents residues favorable to glycosylation, while blue is used to show residues that are unfavorable to glycosylation. The histogram on the right shows average value for amino acids not in the i position. (c) Boxplots of peptide scores in the positive and negative sets, as well as boxplots of peptide scores for individual proteins in the negative set.

The optimized matrix (Fig. 3b) suggests that glycosylated peptides have a bias for the methyl group-containing amino acids alanine, valine, methionine, threonine, isoleucine and leucine; proline was also favorable. Glycosylated peptides were depleted in glycine, glutamine and asparagine as well as the charged amino acids, glutamate, aspartate and arginine. Trends for glycine, alanine, proline, asparagine and glutamine seemed to be consistent across the length of the matrix. The results of the optimization are shown in Figure 3c, which demonstrate that the positive and negative sets are largely separated. Notably, peptide scores for individual IDRs in the negative dataset are quite variable. Peptides derived from low-complexity IDRs that readily phase separate, such as DDX4, FMRP and TAF15, are overrepresented in the negative set due to the bias in protein availability in our lab. The smaller number of peptides derived from CFTR and SARA do not score as poorly as the other peptides in the negative set, indicative of a deficiency in the negative set (see below). The similarities of the matrix values along the length of the peptides suggested that OGT might select IDRs with particular amino acid makeups, rather than exclusively selecting short linear motifs via the active site. We refer to our predictive algorithm that utilizes the compositional bias of 39 residues around the modification site as OGTcomPred.

### Testing Compositional Bias

To measure compositional bias in the positive and negative datasets, we used the program fLPS2.0(65). The background proportions of amino acid types were those derived from human UniProt records(53) or alternatively a dataset of disordered proteins (Fig. S1). To determine the proportions of amino acids in disordered proteins, we determined their abundance in a MobiDB(66) manually curated version of the DisProt database(67). We further selected only human proteins with greater than 50%fractional disorder. The compositional biases were more evident in the PhosphoSite database, due to the larger size of the database, when compared to the experimental negative set (Table 2). As expected, there is a bias for inclusion of serine and threonine in the positive dataset, since these residues are present in every peptide in the database, that was observed when either the human proteome amino acid composition was used or when the disordered protein amino acid composition was used. In contrast, the negative dataset does not appear biased for threonine in either case, suggesting that threonines might be more favorable for glycosylation. As suggested by the substrate scoring matrix, there is a bias for inclusion of methyl-containing residues like alanine and valine and for isoleucine and methionine when using disordered protein amino acid composition. The positive set also has a bias for prolines, though this disappears when using the disordered protein composition. The negative set has a notable bias for inclusion of glycine and to a lesser extent glutamine and asparagine. Thus, analysis of the training datasets support a difference in the compositional biases of the positive and negative datasets and a role for amino acid compositional bias in glycosylation target selection.

**Table 2.** Measurement of compositional bias in the positive and negative datasets. Compositional biases of low probability were assessed for the PhosphoSite database of O-GlcNAc sites (positive set) and our experimentally determined negative set as a whole using the method of Harrison and Gerstein as implemented in fLPS 2.0. The background amino acid composition was either the human proteome composition or the amino acid composition of a set of disordered proteins (see methods).

| Dataset | Residues with Pbias < $10^{-3}$ using human proteome composition |
| --- | --- |
| PhosphoSite database | T (< $10^{-264}$), S (< $10^{-264}$), P ($10^{-264}$), A ($10^{-82}$), V ($10^{-12}$), G($10^{-3}$) |
| Experimental negative set | G ($10^{-16}$), S ($10^{-8}$), Q ($10^{-4}$), R ($10^{-3}$), N ($10^{-3}$) |
| | Residues with Pbias < $1 \times 10^{-3}$ using disordered protein database composition |
| PhosphoSite database | T (< $10^{-264}$), S ($10^{-239}$), V ($10^{-101}$), A ($10^{-51}$), I ($10^{-13}$), F ($10^{-5}$), Y ($10^{-4}$), M ($10^{-4}$) |
| Experimental negative set | G ($10^{-9}$), Y ($10^{-5}$), S ($10^{-4}$), N ($10^{-4}$) |

### Compositional Mutations

To test the effect of amino acid composition on glycosylation, we used the trends from our optimized matrix incorporated into our OGTcomPred algorithm and the compositional bias measures to predict mutations in the FUS LCR_N_ that would enhance glycosylation. We made six constructs containing various combinations of mutations. Mutation were chosen on the basis of composition with no regard for local sequence motifs, in order to test the role of composition rather the role of specific glycosylation motifs. In total, thirteen glycines were mutated to alanine, threonine or proline. Long stretches of alanines were avoided to prevent α-helix formation. The remaining substitutions were glutamines mutated to threonine or proline, serines mutated to threonines and a single aspartate mutated to threonine. Mut-A contained the mutations Q31T, G34A, Q36A, G40T, Q43P, D46T and G49A. Mut-B contained the mutations G67A, Q69T, G74A, G76P, G79A, G80P, G82A, S83T and Q85P. Mut-C contained the mutations G99A, G101T, S107T, S108T, G111A, G114A and S115T. Mut-D combined Mut-A and B mutation. Mut-E combined Mut-B and C mutations. Mut-F combined mutations from Mut-A, B and C (Supplementary Table 1). The total number of serines and threonine increased by less than 10% going from WT to Mut-F. SUMO fusions of the mutants Mut-B through Mut-F were successfully purified, glycosylated and subjected to LC-MS. We failed to purify Mut-A. Unlike in the initial experiment with the three FET proteins (Fig. 1), SUMO fusions with the LCR_N_ were used in the glycosylation reactions and the mass spectrometry experiments as we could not consistently get data without the fusion tag in place. However, the SUMO may have reduced the level of glycosylation possibly via transient steric inhibition. Under the conditions used in this experiment, we observed only a single O-GlcNAc modification on WT FUS. With an increasing number of mutations, higher O-GlcNAc stoichiometries were observed, with as many as seven on the mostly highly mutated construct, Mut-F (Fig. 4a). Comparing the maximum number of observed sites with the number of sites predicted by our OGTcomPred algorithm gave a Pearson correlation of 0.84 with a p value of 0.038 (Fig. 4b). In contrast, comparing the number of sites predicted by O-GlcNAcPredII, considered to be the best existing predictor(45), gave a Pearson correlation of 0.59 with a p value of 0.21 for this set. The data for individual mutants showed a distribution in the number of glycosylation sites, matching expectation. However, for mut-D, peptides with one, three and four added sugars were observed, but peptides with two added sugars were not observed (Fig. 4a). The explanation for this is unclear, though it is possible that this peptide simply was not detected in the mass spectrometer. In summary, the FUS glycosylation mutations support the hypothesis that OGT can utilize amino acid composition over significant stretches within IDRs to recognize substrates and suggests that the relatively short peptide sequences used by O-GlcNAcPredII to identify glycosylation sites do not fully capture this compositional bias.

**Figure 4.**
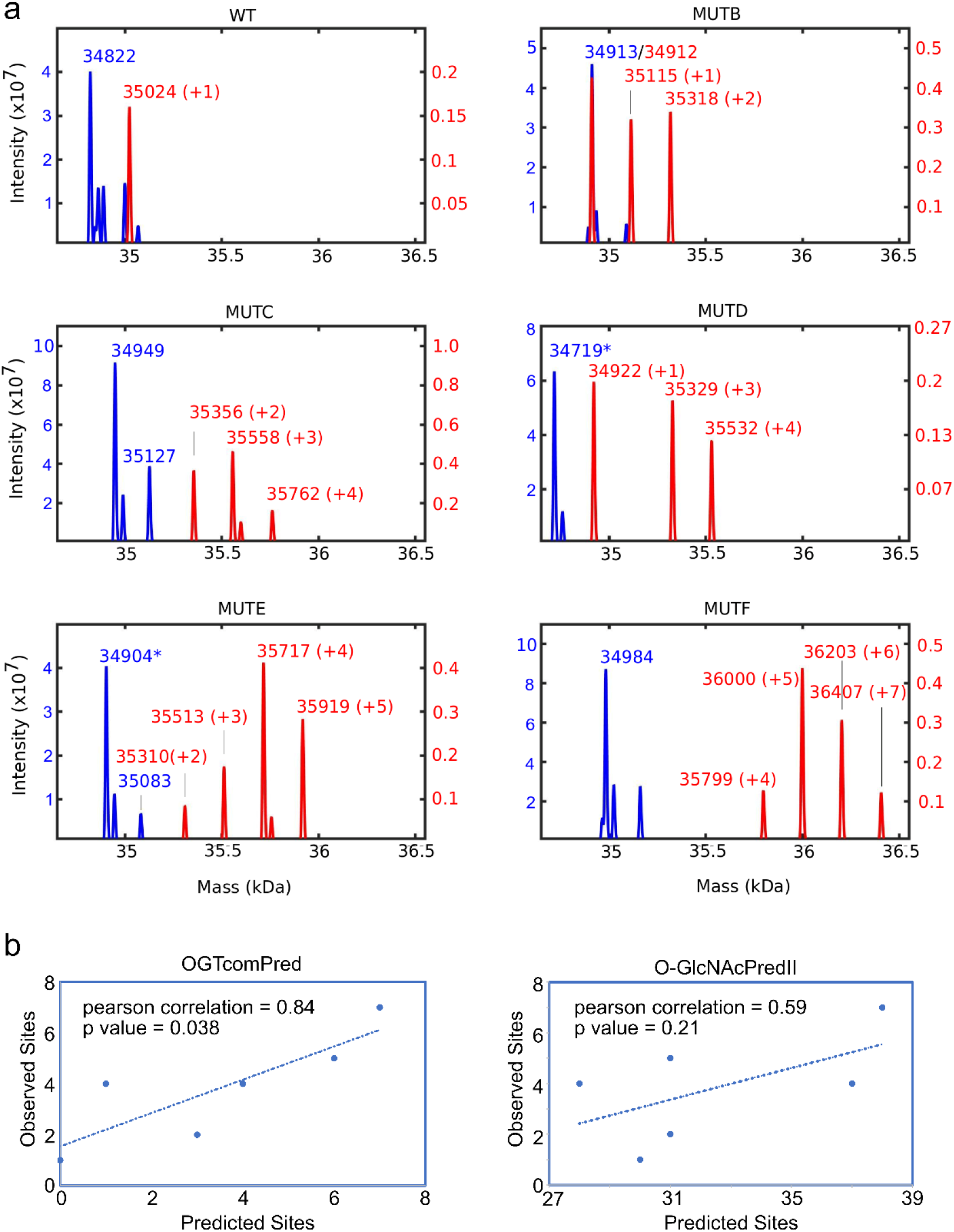
Glycosylation of FUS compositional mutants. (a) Mass spectrum of SUMO fusions of WT and mutant FUS LCR_N_, before (blue) and after (red) overnight glycosylation by OGT. (b) Correlation plots of the maximum number of observed sites versus sites predicted by OGTcomPred or O-GlcNAcPredII.

### NMR evidence for direct interaction between OGT-TPR and FUS

Since it is known that some OGT substrates require the TPR for efficient glycosylation, we hypothesized that the TPR functions by binding to substrates to increase the likelihood of contact with the catalytic domain. NMR is a reliable means of confirming protein interactions involving conformationally flexible IDRs. Therefore, to test whether the TPR can bind to FUS, we generated NMR spectra of ^15^N-labelled WT FUS LCR_N_ in the presence and absence of unlabelled TPR region of OGT (OGT-TPR, aa 2-474) fused to a SUMO tag. The ^15^N labelling allows us to observe spectral peaks (circles in Fig. 5) that correspond to individual amide protons in FUS. In the overlay of the WT FUS spectra with and without OGT-TPR (Fig. 5a), we observe that addition of OGT-TPR causes several peaks to largely disappear, specifically, peaks arising from the two SYXGY motifs (motifs found at aa 37-41 and 96-100) in FUS. In Fig. 5c, a plot of signal intensity ratios with and without OGT-TPR demonstrates the heterogeneity in peak intensity changes, with intensity losses ranging from none to 90% and an average peak intensity ratio of 0.48 +/− 024. The simplest mechanistic explanation is that FUS binds to the OGT-TPR, which is 50 kDa in size and is known to form a dimer, causing the rotational motion of FUS-interacting residues to slow dramatically and leading to significant NMR signal loss. Residues further from the directly interacting residues experience less restriction in rotational motion and consequently less signal loss. The heterogeneous peak intensity loss provides solid evidence that the OGT-TPR binds to WT FUS in a dynamic manner, with multiple interacting elements of FUS exchanging on and off the surface of the TPR(68), and suggests that some parts of the FUS sequence are preferred binding sites. To test whether the compositional mutations enhance binding to OGT-TPR, we next recorded NMR spectra of FUS Mut-F in the presence and absence of OGT-TPR. The Mut-F overlay shows a more dramatic loss of signal intensity in the presence of OGT-TPR (Fig. 5b and 5d), with an average peak intensity ratio of 0.36 +/− 0.22, strongly suggesting that the compositional mutations enhance binding to the OGT-TPR. To control for possible binding of SUMO to WT FUS and FUS Mut-F, we repeated the experiment using OGT-TPR not fused to SUMO. The results were qualitatively similar (Fig. S2) providing evidence that changes in the FUS spectra are due to TPR binding and not SUMO binding. However, in the absence of the SUMO, the samples with OGT-TPR phase separated which made quantitative comparison of the apo and plus OGT-TPR samples impossible.

**Figure 5.**
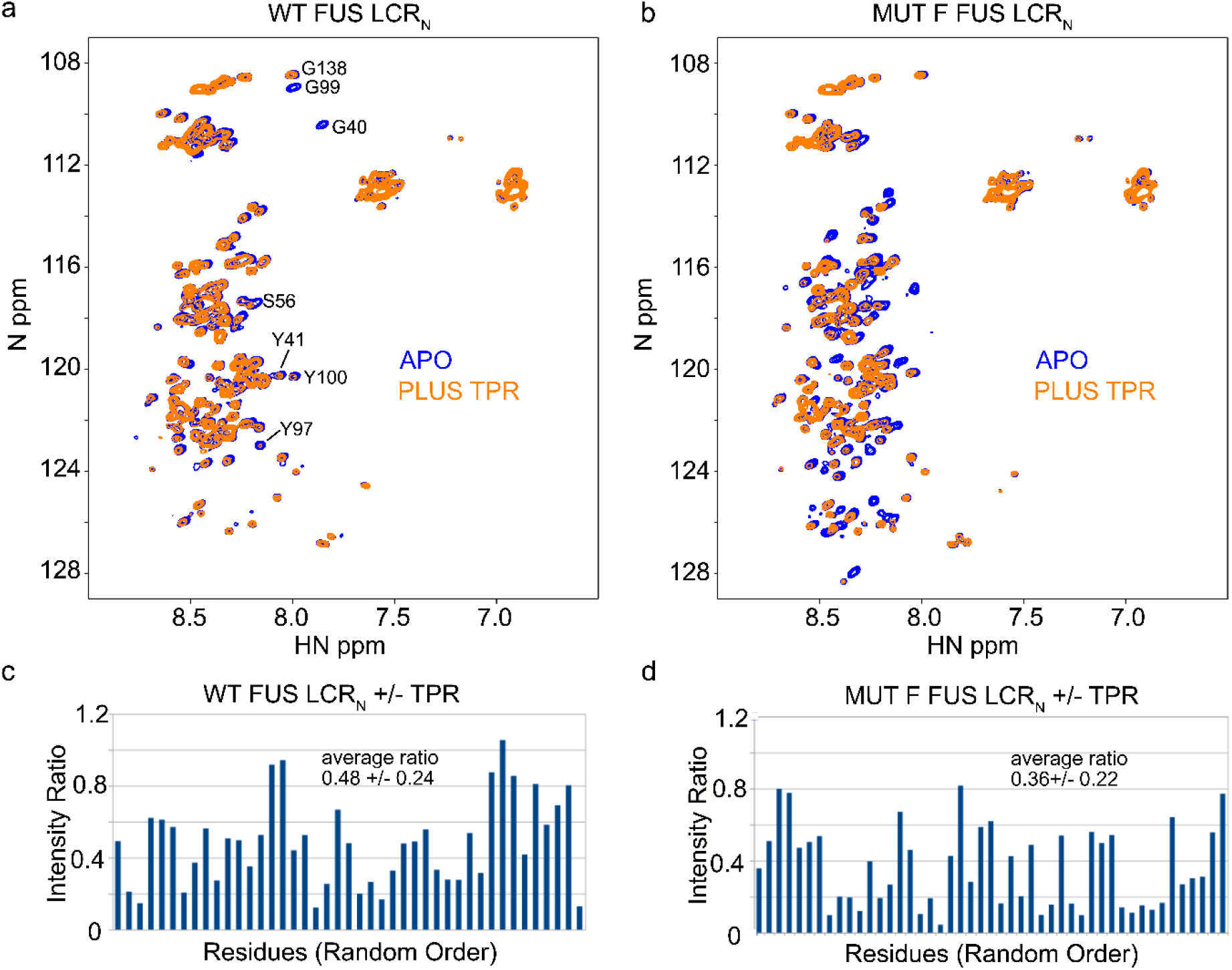
OGT-TPR interaction with FUS and Mut-F FUS. ^1^H-^15^N HSQC spectra of WT FUS LCR_N_ (a) and Mut-F FUS LCR_N_ (b) in the presence and absence of SUMO-OGT-TPR. Spectra of the FUS LCR_N_ at 20 μM are shown in blue and spectra in the presence of 32 μM SUMO-OGT-TPR are shown in orange. Spectra were recorded with a field strength of 600 MHz at 5°C in a buffer comprised of 40 mM KPO_4_, 125 mM NaCl, 0.5 mM EDTA, 0.5 mM benzamidine, 5 mM DTT and 10% D_2_O, pH 7.2. Resonance assignments for some peaks in the WT spectrum were obtained by transferring some assignments from BMRB record 26672. SUMO-control experiments are shown in Figure S1. Plots (c) and (d) show intensity ratios (plus OGT-TPR/apo) for WT and Mut-F FUS LCR_N_ respectively. Intensity ratios are shown in random order, since most of the residues were not assigned (see Experimental Procedures).

### Test Case: CREB-binding protein (CBP)

We next tested our compositional matrix glycosylation site predictor, OGTcomPred, on four known IDRs from human CBP (69–71) and then measured glycosylation experimentally using mass spectrometry. The four regions were the ID1 (CBP aa 1-344), ID3 (CBP aa 676-1080), ID4 (CBP aa 1851-2057) and ID5 (CBP aa 2124-2442). We predicted 13, 34, 15 and 6 sites in ID1, ID3, ID4 and ID5, respectively (Fig. 6), whereas only one site in each of ID1 and ID5 and no sites in ID3 and ID4 are listed in the PhosphoSite O-GlcNAc database. Following overnight glycosylation, mass spectrometry demonstrated glycosylation at a median of 3 sites in ID1 and a median of 2 sites in ID5 (Fig. 6). ID4 was predominantly unglycosylated, which could be due to secondary structure elements unaccounted for by the prediction (see Discussion). We were unable to obtain mass spectrometry data on the full ID3, so we digested the glycosylated protein with trypsin and submitted the sample to MS/MS (Table 3). We found glycosylation at S709, with a further three sites between residues 715 and 768 and one between residues 972 and 998. Therefore, there are at least 5 possible glycosylation sites in ID3. These results confirm that OGT can glycosylate more sites than are listed in the PhosphoSite database, but indicates that our predictor shares a high false positive rate with previously developed predictors.

**Figure 6.**
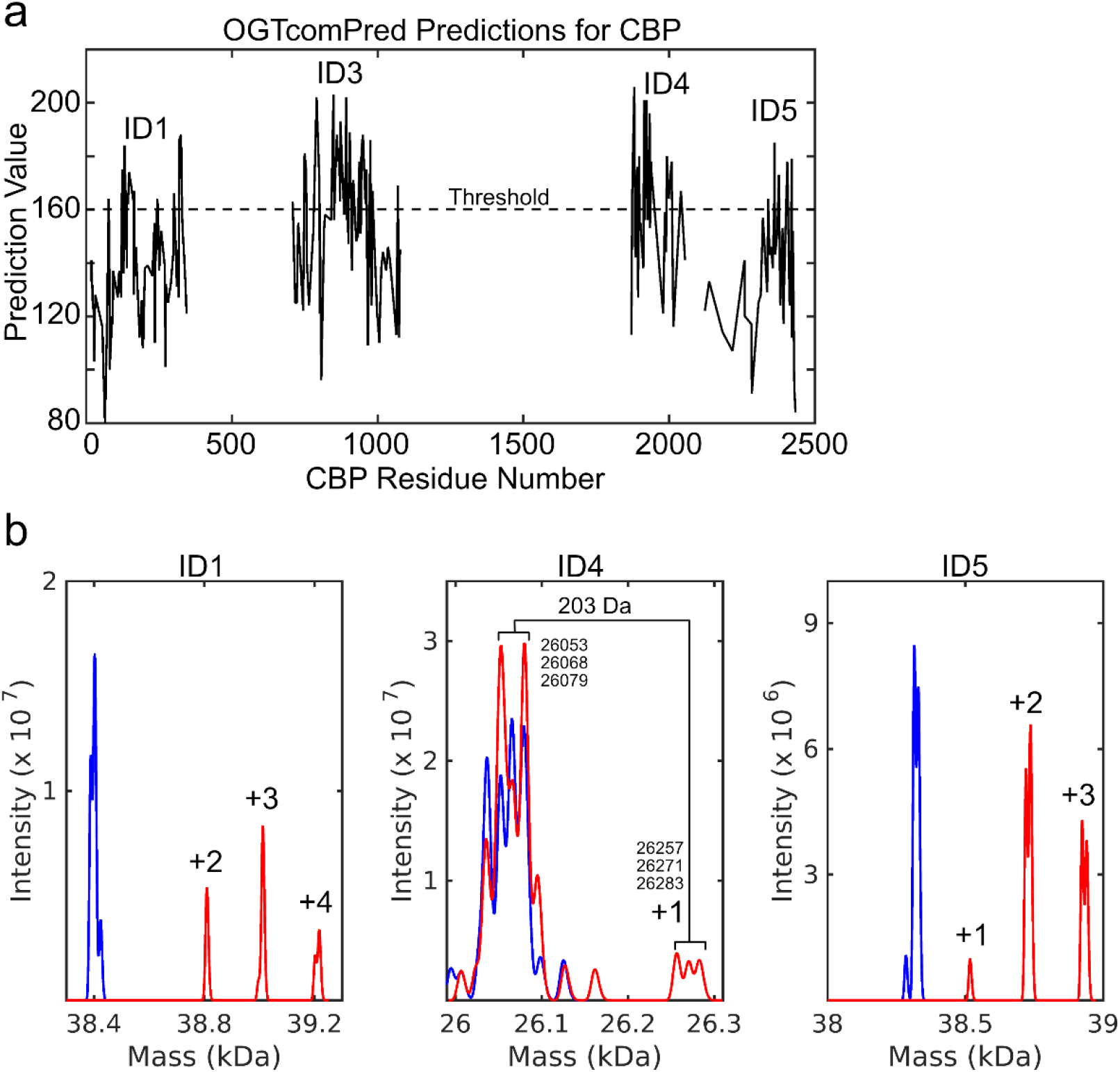
CREB-binding protein (CBP) glycosylation. (a) OGTcomPred prediction of OGT glycosylation sites for 4 intrinsically disordered segments of CBP. (b) Mass spectrum of CBP ID1, ID4 and ID5 before (blue) and after (red) glycosylation. Peaks annotated as +1, +2, +3 and +4 are 203, 406, 609 or 1212 Da bigger than the unglycosylated material. Some of the peaks appear as doublets (ID1 and ID5) because a large fraction of the protein is modified by a single oxidation event (+16 Da). ID4 is modified by several small chemical modifications in addition to the sugar modification (+203 Da).

**Table 3.** Results of MS-MS experiments on CBP ID3. Peptides identified by the MS-MS sequence are listed. The number of glycosylation sites for each peptide was determined by the difference in the molecular weight of the parent ion. One site was identified unambiguously, with a further four sites for which the precise modification sites could not be determined. LC-MS/MS data: DOI: 10.5281/zenodo.6986306.

| CBP ID3 peptide span | Observed glycosylation sites<br>(total number of serines and<br>threonines in peptide) | Confirmed Sites |
| --- | --- | --- |
| 676-714 | 1(1) | S709 |
| 715-742 | 2(3) |  |
| 743-768 | 1(6) |  |
| 972-998 | 1(6) |  |

### Dataset Analysis

To gain further insight into OGT substrate recognition, we analyzed matrix plots of position-dependent amino acid frequencies for the positive and negative datasets used here and in the O-GlcNAcPRED-II predictor development (Fig. 7). The O-GlcNAcPRED-II positive dataset (Fig. 7b) and the PhosphoSite dataset (Fig. 7a) from which it is derived are highly similar. In contrast, the experimental negative set from this study (Fig. 7c) and the negative set for the O-GlcNAcPRED-II study (Fig. 7d) are quite different. Although published details on how the O-GlcNAcPRED-II negative dataset were obtained are limited, it contains approximately 51,000 peptide sequences. This database is large and, at first glance, appears to have a very limited amount of residual sequence-specific information with nearly uniform amino acid frequencies along the length of the peptide. As such the database may have been useful as a way to normalize the positive dataset against expected amino acid frequencies, rather than primarily contributing information on sites that are difficult to glycosylate. Consistent with this, the O-GlcNAcPRED-II negative dataset matrix is very similar to the matrix for all human protein S/T centred peptides derived from UniProt (Fig. 7e). Examination of the positive datasets demonstrates overall amino acid frequencies that are similar to the O-GlcNAcPRED-II negative set and the human proteome. For example, serines and to a lesser extent prolines, alanines, glycines and leucines are present with high frequency in the positive datasets and the O-GlcNAcPRED-II negative dataset. However, in the positive sets, one also sees amino acids that are over- or under-represented in a position-dependent manner relative to the O-GlcNAcPRED-II negative set. These primarily occur within the 4 residues before and after the serine/threonine glycosylation site and likely represent sequence specific elements that are recognized by the catalytic domain of OGT. For example, peptides with prolines in the i-3, i-2 and i+2 position seem to be favorably selected by OGT. The presence of many threonines in the i+1 through i+14 indicates a preference for threonines C-terminal to the serine/threonine glycosylation site, an observation that has previously been reported(72). In contrast to the O-GlcNAcPRED-II negative set, the experimental negative set presented here is extremely small, just 135 peptides with overlapping sequences, and is likely not a very good sampling of the OGT negative site proteome. The small size of the dataset results in a rather noisy dataset. Nonetheless, the amino acid frequencies differ significantly from the human proteome as shown by the compositional bias results above. In the matrix representation of the negative set, glycine and to a lesser extent arginine, aspartate, asparagine and glutamine have a higher relative abundance compared to the other datasets. At the same time alanine, proline, valine, isoleucine, leucine and methionine are less abundant than in the other datasets. While the small size and biased nature of the negative dataset make strong conclusions unwise, the compositional bias is suggestive. Of note, the noise in the negative dataset precludes discernment of any sequence specific information.

**Figure 7.**
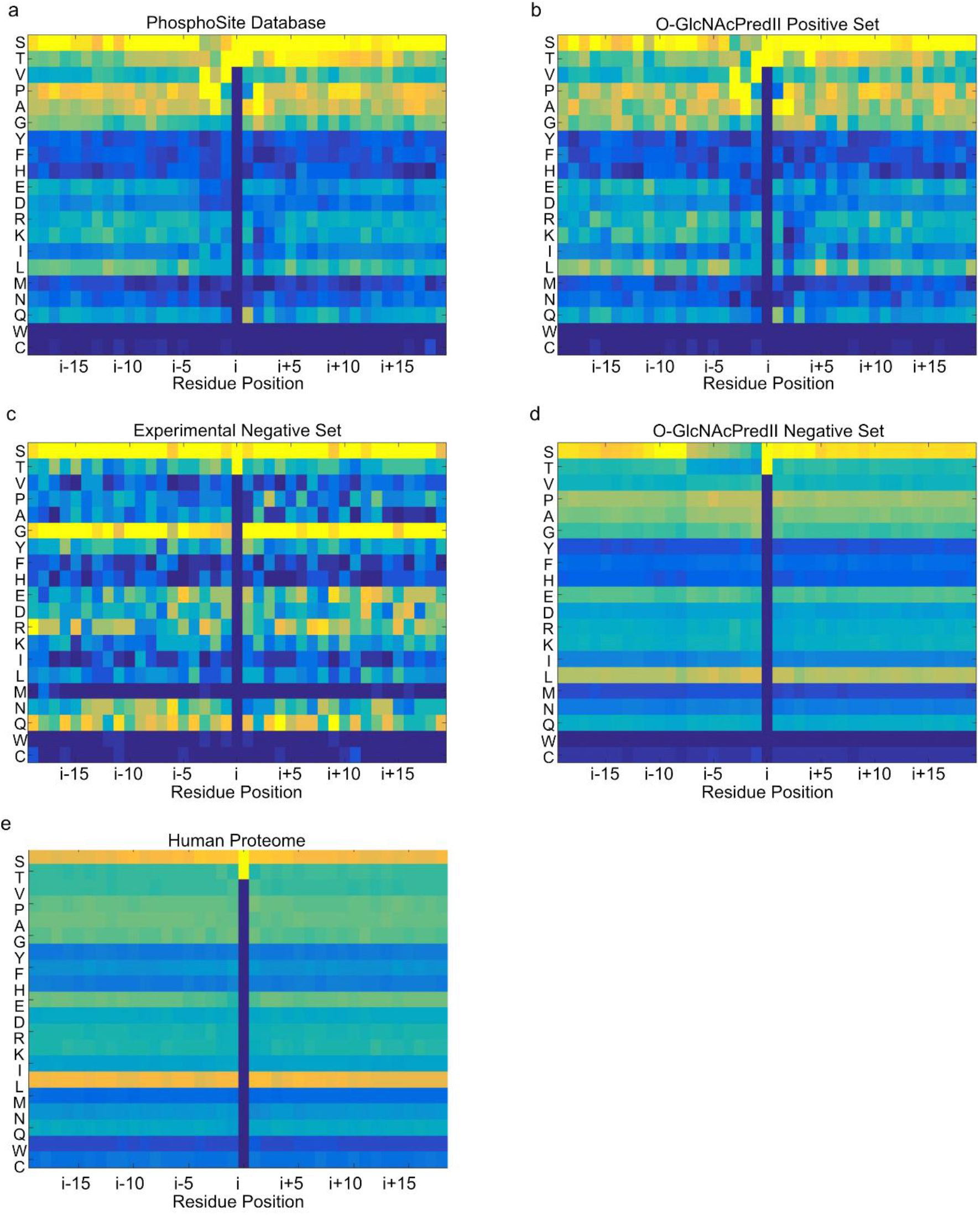
Prediction Dataset Analysis. Positional amino acid frequency plots for the (a) PhosphoSite Database of O-GlcNAc sites, (b) O-GlcNAcPredII positive dataset, (c) the experimental negative dataset developed here, (d) the negative dataset used in the development of the O-GlcNAcPredII predictor, and (e) the human proteome (UniProt database). Residue positions are relative to the serine or threonine at the glycosylation site. Frequency is indicated by a blue-yellow gradient, with yellow representing high frequency and blue indicating low frequency.

### Comparison to other O-GlcNAcylation predictors

Rigorous comparison of site predictors requires definitive knowledge of both positive and negative sites, since specificity and accuracy cannot be calculated without knowing the number of negative sites. Our knowledge of negative sites is still extremely limited, in part due to a focus on high-throughput approaches, which are better at identifying positive sites. Lamin A is an O-GlcNAcylation target that has been studied in a targeted low-throughput approach, giving more confidence that sites not identified as glycosylated are in fact not glycosylated by OGT(47). We used Lamin A as a test case to crudely compare the different predictors (Table 4). Results from our simple predictor OGTcomPred compare favorably with early predictors, though they are not as good as more sophisticated tools such as O-GlcNAcPRED-II(41)). Interestingly, preliminary exploration suggests that adding some sequence specificity back into our predictor improves prediction results while decreasing the ability of our predictor to discriminate between our positive and negative datasets (not shown). This supports our suspicion that some sequence specificity is lost due to the small size of our experimental negative set. Nevertheless, the fact that our predictor OGTcomPred compares well with some of the other predictors, despite this loss, supports our contention that OGT substrates have a compositional bias.

**Table 4.**
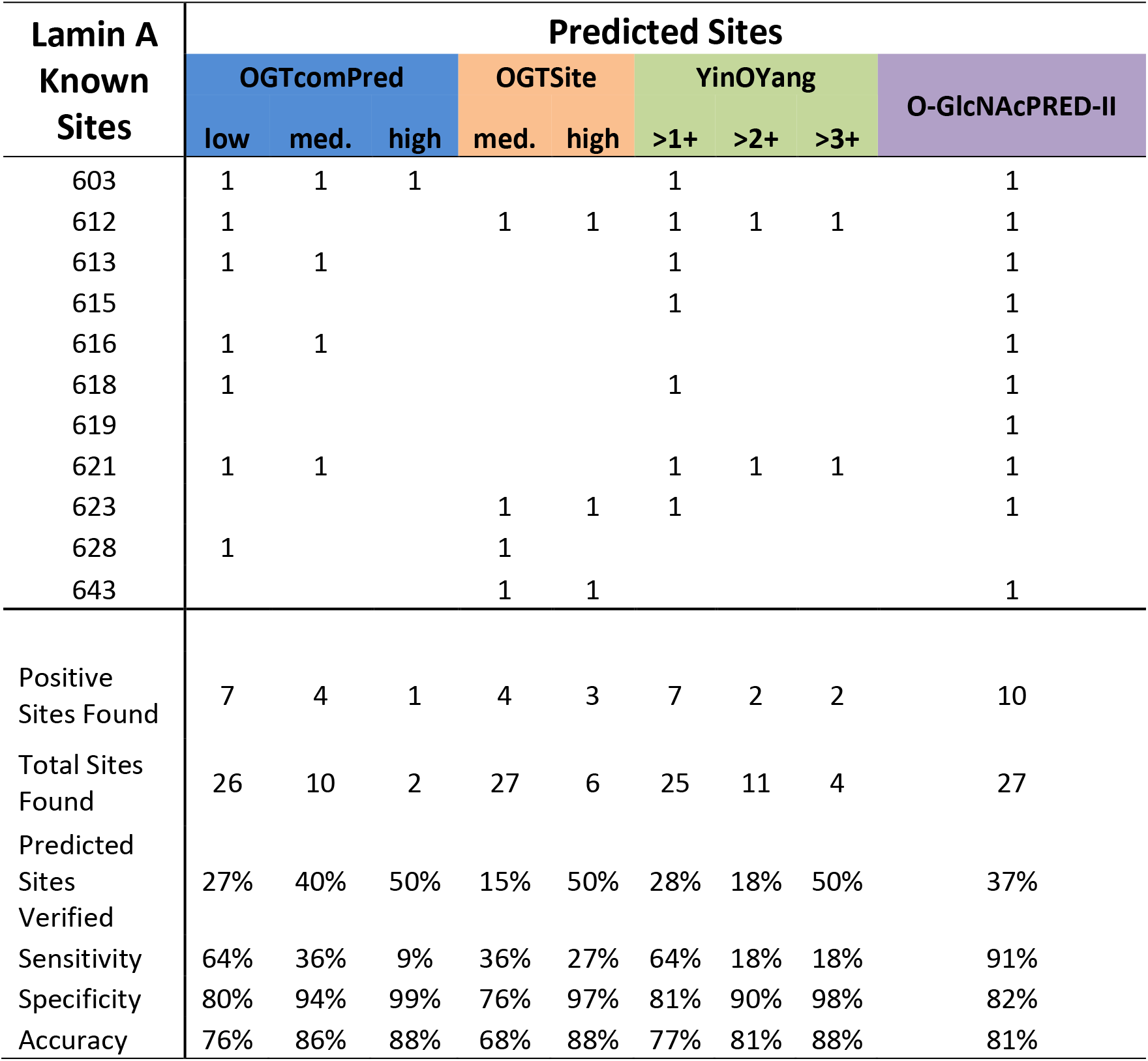
Assessment of Sensitivity, Specificity and Accuracy for O-GlcNAc predictors using Lamin A as a test case. Known sites and prediction values for these sites are listed in the top half of the table and used to assess sensitivity, specificity and accuracy in the bottom half of the table. Assessments were performed at different threshold levels when possible.

| Lamin A<br>Known<br>Sites | Predicted Sites |  |  |  |  |  |  |  | O-GlcNAcPRED-II |
| --- | --- | --- | --- | --- | --- | --- | --- | --- | --- |
|  | OGTcomPred |  |  | OGTSite |  | YinOYang |  |  |  |
|  | low | med. | high | med. | high | >1+ | >2+ | >3+ |  |
| 603 | 1 | 1 | 1 |  |  | 1 |  |  | 1 |
| 612 | 1 |  |  | 1 | 1 | 1 | 1 | 1 | 1 |
| 613 | 1 | 1 |  |  |  | 1 |  |  | 1 |
| 615 |  |  |  |  |  | 1 |  |  | 1 |
| 616 | 1 | 1 |  |  |  |  |  |  | 1 |
| 618 | 1 |  |  |  |  | 1 |  |  | 1 |
| 619 |  |  |  |  |  |  |  |  | 1 |
| 621 | 1 | 1 |  |  |  | 1 | 1 | 1 | 1 |
| 623 |  |  |  | 1 | 1 | 1 |  |  | 1 |
| 628 | 1 |  |  | 1 |  |  |  |  |  |
| 643 |  |  |  | 1 | 1 |  |  |  | 1 |
| Positive<br>Sites Found | 7 | 4 | 1 | 4 | 3 | 7 | 2 | 2 | 10 |
| Total Sites<br>Found | 26 | 10 | 2 | 27 | 6 | 25 | 11 | 4 | 27 |
| Predicted<br>Sites<br>Verified | 27% | 40% | 50% | 15% | 50% | 28% | 18% | 50% | 37% |
| Sensitivity | 64% | 36% | 9% | 36% | 27% | 64% | 18% | 18% | 91% |
| Specificity | 80% | 94% | 99% | 76% | 97% | 81% | 90% | 98% | 82% |
| Accuracy | 76% | 86% | 88% | 68% | 88% | 77% | 81% | 88% | 81% |

## Discussion

It is not yet possible to reliably predict O-GlcNAcylation sites despite there being a significant amount of effort put towards developing predictors for O-GlcNAcylation sites. Here we developed a small, experimentally-tested negative dataset, which suggests that OGT has the ability to distinguish between substrates and non-substrates based on amino acid composition over an extended sequence length, a factor that should be a consideration in predicting glycosylation. Specifically, methyl group-containing amino acids and proline were favorable, while glycine, glutamine and asparagine as well as the charged amino acids, glutamate, aspartate and arginine seem to inhibit glycosylation. Compositional mutations introduced into the FUS LCR_N_ with no regard for specific active-site recognition motifs support this idea. Changes in glycosylation stoichiometry were correlated with the number of glycosylation sites predicted by our compositional matrix predictor OGTcomPred. NMR data provide evidence that glycosylation-promoting compositional mutations enhance OGT-TPR binding to the FUS LCR_N_. Together, these observations support a model in which interactions between intrinsically disordered substrates and the OGT-TPR can facilitate glycosylation of sites that have a suboptimal interaction with the catalytic site, as has been previously observed(73).

There are many factors that affect OGT recognition *in vivo*. Here we focused on factors that influence the ability of ncOGT (OGT with 13 TPR repeats) to directly recognize and glycosylate other proteins *in vitro,* including interactions of the substrate with the catalytic site and the TPR (Fig. 8). While some substrates seem to require the full TPR for efficient glycoslyation, others can be glycosylated with minimal TPR repeats. An example of the latter is a 12-amino acid substrate peptide derived from the casein kinase II (CKII), which can be glycosylated by a shortened OGT variant that is missing 5.5 TPR repeats relative to the full ncOCT variant(31). Furthermore, addition of TPR *in trans* does not competitively inhibit glycosylation of CKII(31), suggesting that the TPR region does not contribute significantly to recognition of CKII as a substrate. In contrast, other substrates like TRAK1(31) and the C-terminal domain of RNA polymerase II(30) require all of the TPR repeats for efficient glycosylation. OGT with a full TPR is known to glycosylate a broader range of substrates than OGT isoforms with fewer TPR repeats(26). This is consistent with OGT requiring a threshold level of affinity for efficient glycosylation, with that affinity being achieved either by optimal interaction between a short peptide segment of an IDR and the OGT catalytic region or alternatively by a combination of many weak interactions between a long IDR and the OGT catalytic region and the TPR.

**Figure 8.**
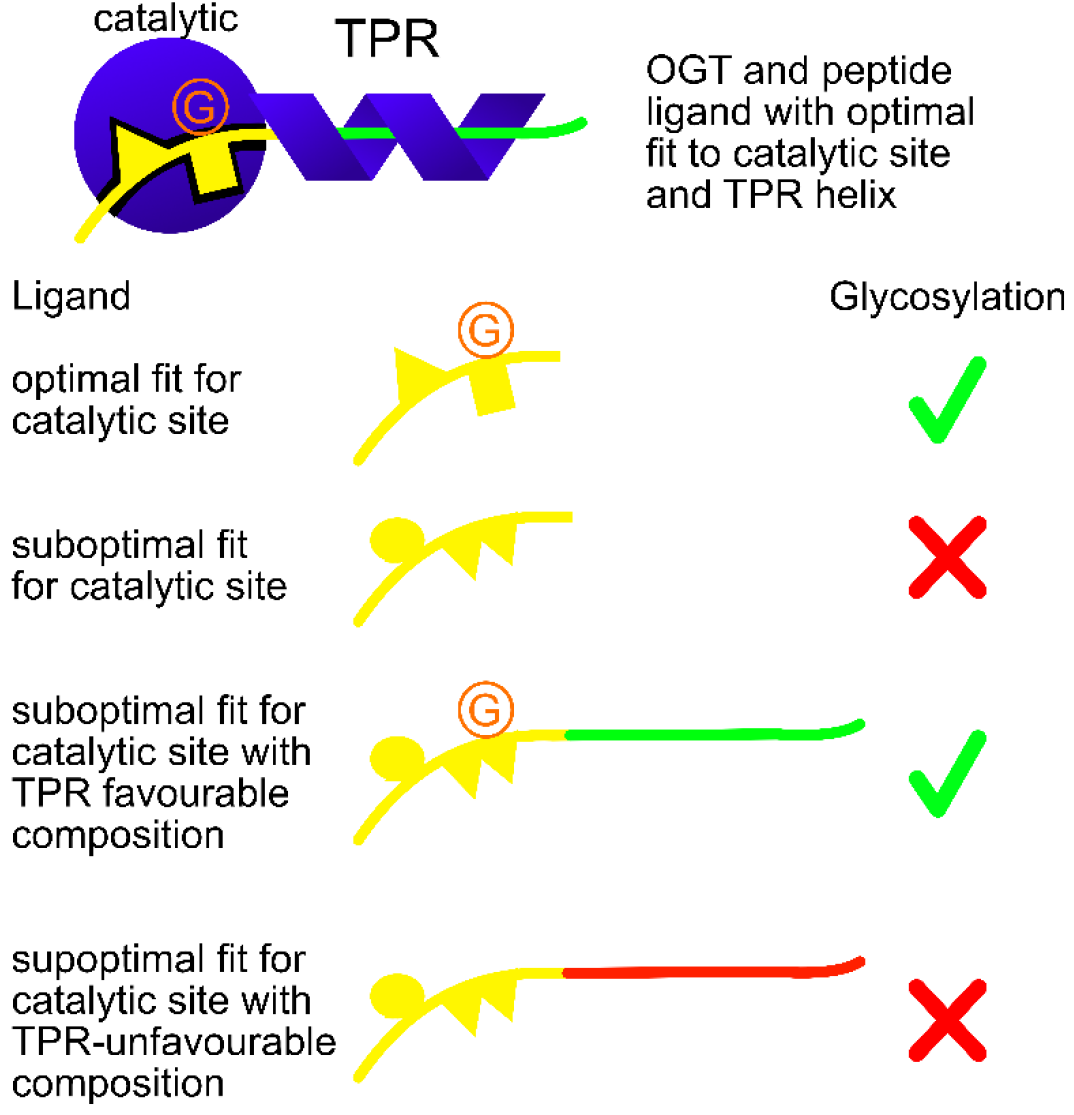
OGT ligand selection model. The OGT catalytic domain and TPR helix are shown in blue, bound to a peptide ligand that has an optimal fit to the catalytic site and a TPR interacting region with a compositional bias that promotes interaction with the TPR. Glycosylation is indicated by the orange G. Short peptides with optimal fits for the catalytic site can be glycosylated, but short peptides with suboptimal fits are not glycosylated. Extended peptides with suboptimal catalytic site fit can still be glycosylated if the peptide has a compositional bias that is suitable for TPR interaction (green), but not if the compositional bias is less favorable for a TPR interaction (red).

### Catalytic Site Interactions

From a structural perspective, it is not yet well understood how substrates are recognized by the catalytic site. Consistent with the wide array of OGT substrates, there are relatively few contacts between the catalytic site and the sidechains of peptide substrates, with crystal structures demonstrating that most contacts involved the backbone of substrate peptides (2, 22). Nonetheless, a screen of randomly-generated 13-residue peptides shows a highly specific selection of substrates for this class of short peptide, with less than 10% of the peptides in the screen being glycosylated as efficiently as the positive control. Crystal structures of multiple substrates demonstrate a highly constrained peptide backbone in the −3 to +2 region involving hydrogen bonds to the peptide backbone. These structures suggest the presence of size preferences or steric restrictions in the different positions along the substrate. For example, smaller amino acids are preferred in the −3 and +2 position, while the −2 position seems to disfavor small amino acids such as alanine and glycine. Even dramatic substitutions of single residues, for example, replacing serines and alanines in the +2 position with phenylalanine, merely reduce the efficiency of glycosylation, introducing energetic costs that might be overcome through TPR interactions with longer peptides. However, the additive effects of several unfavorable changes could possibly prevent glycosylation. So, rather than trying to define sequences that interact optimally with the catalytic site, it might be more helpful to look for sequences that prohibit glycosylation. The large number of possible sequence combinations will make uncovering prohibitive sequences difficult, but may be a key piece of solving the prediction puzzle.

A second piece of the puzzle is trying to define the amino acid preferences for interactions with the OGT-TPR. In the crystal structure of OGT-TPR bound to a peptide derived from HCF-1, the substrate is in an extended conformation in the inside of the helix. A series of conserved asparagine residues arranged on the inside of the TPR helix(24) form bidentate interactions with the substrate peptide backbone(72). Mutation of five of these asparagines selectively inhibited a substantial number of substrates that require the OGT-TPR for efficient glycosylation(52). Since these are backbone contacts, they likely play a minimal role in specificity. Structures with an HCF-1-derived peptide bound inside the helix also show four TPR aspartates forming hydrogen bonds with threonine sidechains in the HCF-1 peptide. This explains the prevalence of threonines C-terminal to the glycosylation site in the Phosphosite and O-GlcNAcPRED-II positive datasets. Our work suggests that glycines are unfavorable, possibly because they are less conformationally restricted, which would impose a greater entropic cost for binding to the asparagine ladder. We also found that small hydrophobic residues such as alanine, valine and proline are favorable. We speculate that these can make favorable van der Waals interactions with the concave surface of the TPR helix, possibly via transient, dynamic interactions(64) that allow substrate sidechains to be correctly positioned in the catalytic site. In contrast, amino acids with sidechains that can form hydrogen bonds seem unfavorable with the exception of threonines and serines. Hydrogen bond-forming amino acids such as asparagine, glutamine and glutamate may introduce geometric constraints that are difficult to satisfy. Glutamate may additionally be unfavorable because of the net excess of acidic residues on the concave surface of the TPR. Consistent with this, OGT constructs with fewer TPR repeats more readily glycosylate substrates with polar uncharged and charged residues such as glutamine, asparagine, lysine, glutamate and aspartate(26).

### Substrate structure impact on glycosylation

Of note, based on existing OGT crystal structures, secondary structure elements and folded domains are predicted to be incompatible with glycosylation by OGT due to steric clashes(22). This could explain our observation that ID4 of CBP does not get glycosylated despite our prediction. In fact, the regions that flank the majority of our predicted target serines and threonines in this intrinsically disordered segment form alpha-helical structures with significant populations(70). We speculate that disruption of these secondary structure elements could promote glycosylation of ID4. Amino acids conducive to maintaining an extended or random coil structure significantly affect the propensity of a peptide to be glycosylated and may explain the preference for prolines and beta-branched residues(2). Although the vast majority of OGT substrates are in IDRs(5), there are a few examples of proteins that are glycosylated in folded or ordered regions, including HBGB-1(74), H2B(75) and αB-Crystallin(76). Glycosylation of these ordered sites could occur if the TPR is able to move away from the catalytic site, as has been suggested by a recent electron microscopy structure(77). Alternatively, the OGT-TPR might be able to unfold a select group of ordered regions, as the TPR-containing karyopherin proteins are known to behave as chaperones(78). Finally, glycosylation of these ordered regions could occur co-translationally before the proteins are fully folded(79, 80). Since most OGT ligands are IDRs, we intentionally picked IDRs to build our negative dataset. When making predictions for proteins for which the structure is unknown, we also couple the prediction to a disorder predictor. Attaining more accurate predictions will likely require incorporation of structural and steric constraints, which may be facilitated by recent advances in structure prediction(81, 82).

How proximal structured elements impact glycosylation is not yet well defined. The range of possible OGT-TPR entry points and the effect of adjacent folded domains on TPR entry are unknown. Examining the TPR structure, it appears that peptides do not need to enter the TPR helix from one end, since there is sufficient space for a peptide to enter the TPR interior from points along the helix. However, larger structural elements would not be able to enter the interior without significant rearrangement of the helix. Thus, our observation that the isolated IDR3 of CBP can be glycosylated *in vitro* does not mean that it can be glycosylated *in vivo* since it is flanked by folded regions in the full-length CBP protein. These considerations add further complexity to prediction efforts. Prediction approaches to date have taken a structure-agnostic approach, but pushing predictions towards higher accuracy will require addressing these structural issues. Overcoming this complexity is a worthwhile goal given the importance of O-GlcNAc modification for modulating protein thermodynamics, aggregation and phase separation propensity.

## Experimental Procedures

### Expression and Purification of Proteins

All DNA constructs were verified by sequencing. Proteins were expressed in *E. coli* BL21 (DE3) RIPL cells using LB media, unless otherwise stated. Cell cultures were grown to an optical density of 0.8 and then induced with 0.5 mM IPTG and harvested after 16 hours at 18 °C. Purifications were carried out at room temperature unless otherwise stated. Purified protein samples were further verified by mass spectrometry to ensure that they were the expected molecular weight.

#### EWS, FUS and TAF15 LCR_N_ purification

His-tagged SUMO fusions of LCR_N_ fragments of human EWS (aa 1-264), FUS (aa 1-214), and TAF15 (aa 1-210) were lysed by sonication and then purified by nickel affinity chromatography using a buffer containing 20 mM CAPS, pH 11, 500 mM NaCl, 4 M guanidinium chloride (GdmCl) with 20 mM imidazole added to the aliquot used for lysis and washing and 280 mM imidazole used in the elution aliquot. Proteins were then subjected to size exclusion chromatography using a buffer comprised of 40 mM arginine, pH 9. A HiLoad Superdex75 HR 16/600 column (Cytiva) was used for all of the size exclusion chromatography described here. Only the purest fractions were retained for glycosylation reactions and mass spectrometry. ULP1 protease (purified in-house) was used to remove the His-SUMO fusion protein. The LCR_N_ protein was then loaded onto a size exclusion column without first concentrating the protein, since concentrating the protein led to significant loss. The same 40 mM arginine pH 9 buffer was used for this step.

#### FUS and FUS mutant LCR_N_ purification

We modified the purification to more reliably obtain FUS or mutant FUS LCR_N_ without the SUMO fusion tag. Nickel affinity chromatography and size exclusion chromatography were followed by consecutive purification on a HiTrap Q column (Cytiva) and a 8 ml phenyl Superose column(Cytiva) using buffer with 40 mM arginine, pH 9 and gradients from 50 mM to 1M NaCl and 1 M to 0 M NaCl, respectively. Following cleavage with ULP1 the protein was again purified by phenyl Superose using the same gradient, to yield highly pure FUS LCR_N_.

#### CBP ID1, ID3, ID4 and ID5 purification

Human CBP IDRs (ID1 (1-344), ID3 (676-1080), ID4 (1851-2057) and ID5 (2124-2442) were lysed in a buffer containing 6 M GdmCl, 50 mM Tris-HCl, pH 8, 500 mM NaCl and 20 mM imidazole and then loaded onto nickel affinity resin. The protein was eluted from the column using 4 M GdmCl, 50 mM Tris, pH 8, 200 mM NaCl and 500 mM imidazole. The protein was then concentrated and purified by gel filtration chromatography using a buffer containing 20 mM KPO_4_, pH 6.5, 100 mM NaCl and 50 μM EDTA.

#### OGT purification

Following expression of human ncOGT (full length, 1-1046) in *E.coli,* cells were resuspended in a buffer containing 25 mM imidazole, 10 % glycerol, 250 mM NaCl and 25 mM HEPES, pH 7.5, 5 mM β-mercaptoethanol. DNaseI and an EDTA-free protease inhibitor tablet (Sigma) were also added to the lysis buffer. Following lysis by sonication and French press, the protein was purified by nickel affinity chromatography and eluted in the same buffer with 250 mM instead of 25 mM imidazole. Fractions containing pure protein were then dialyzed in 25 mM HEPES, pH 7.5, 40 mM NaCl, 0.5 mM EDTA and 5 mM β-mercaptoethanol and then loaded onto a 5 ml HiTrap Q-XL column (Cytiva) and purified at 4 °C using a gradient from 0.05 to 1.0 M NaCl. Although the protein appeared pure after this anion exchange step, we further purified the protein using a HiLoad Superdex 200 HR16/600 size exclusion column using a buffer containing 40 mM KPO_4_, pH 7.5, 125 mM NaCl, 0.5 mM EDTA, 0.5 mM benzamidine and 5 mM β-mercaptoethanol to ensure that no contaminating proteases remained.

#### OGT-TPR purification

A construct containing SUMO fused to residues 2-474 of ncOGT representing the TPR region was purified by Ni affinity chromatography as for the full-length ncOGT purification. The SUMO tag was cleaved off using ULP. NaCl was then added to the sample to bring the total NaCl concentration up to 1M. This was followed by purification on a phenyl Superose column in a buffer of 25 mM Hepes, pH 7.5, 5 mM β-mercaptoethanol, using a 1 M to 150 mM NaCl. As a final purification step, the OGT-TPR was subjected to size-exclusion chromatography using a buffer of 44 mM KPO4, 137.5 mM NaCl, 0.55 mM EDTA and 0.55 mM benzamidine, pH 7.2. Purification of the OGT-TPR with the SUMO tag cleaved off was similar.

#### Purification of negative set proteins

Human SARA (aa 766-822)(60), DDX4 (aa 1-236)(61), CFTR (aa 654-838)(62) and FMRP (aa 445-632)(63) were expressed and purified as described previously.

#### Production of protein for NMR spectroscopy

Isotopically labelled proteins for NMR spectroscopy were expressed in M9 minimal media using ^15^N ammonium chloride as the sole source of nitrogen.

### OGT reaction conditions

Protein samples were dialyzed into 40 mM K_2_HP0_4_, pH 7.5, 125 mM NaCl, 0.5 mM EDTA, 2 mM betamercaptoethanol and 0.5 mM benzamidine. Reactions were performed at a protein concentration of 20 μM. Following addition of 1 μM ncOGT and 1 mM UDP-GlcNAc, samples were incubated at room temperature for 16 hours.

### Mass Spectrometry

MS experiments were carried out at the Structural Genomics Consortium Toronto facility or at The Hospital for Sick Children SPARC Molecular Analysis facility. Samples were prepared by adding formic acid to a final concentration of 0.1% v/v. To determine glycosylation stoichiometry, purified glycosylated proteins and controls were either run on a Thermo-Fisher Orbitrap Q Exactive High Field instrument or on an Agilent UPLC-quadrupolar time-of-flight (Q-ToF) 6545 MS system equipped with a Dual JS electrospray ionization source. Samples were desalted online via a C18 column. Raw data were either processed using Agilent MassHunter software or Thermo-Fisher software and deconvoluted using the maximum entropy algorithm with appropriate mass ranges. The deconvoluted data were then plotted using MATLAB. To identify specific glycosylation sites in the EWS LCR_N_ region or the ID3 region of CBP, the protein was digested with chymotrypsin or trypsin respectively, and then subjected to LC-MS/MS on a Thermo-Fisher Orbitrap Q-Exactive mass spectrometer, using higher energy collisional dissociation (HCD). To identify glycosylation sites with confidence, we set the following thresholds: the parent ion error had to be less than 1 ppm and the number of fragment ions with a score of less than 7 ppm had to be greater than 8. As the O-GlcNAc modification was lost during the peptide fragmentation step, we were able to identify peptides that were glycosylated (parent ion had modification), but typically unable to identify exactly which residues were glycosylation sites. The MS/MS data were analyzed manually, since the software modification site assignment process assumed that the sugar was still present following the fragmentation step.

### NMR Spectroscopy

HSQC experiments(83) were performed at 5 °C in a buffer containing 40 mM KPO_4_, pH 7.2, 125 mM NaCl, 0.4 mM EDTA, 0.5 mM benzamidine, 5 mM DTT and 10% D_2_O. Matched samples were recorded on 20 μM ^15^N labelled samples (below the threshold for phase separation) of either WT FUS LCR_N_ or Mut-F FUS LCR_N_ in the absence and presence of 32 μM SUMO-fused OGT-TPR. Spectra were processed with NMRPipe(84) and displayed in CCPNMR(85) software. Peak intensities were obtained using Sparky(86) software. Peak assignments for FUS LCR_N_ were obtained from the BMRB(87, 88). However, since our sample conditions differed from the conditions used by Burke et al. (BMRB 26672), only peaks in less crowded regions of the spectrum could be assigned. The experiment was repeated using OGT-TPR with the SUMO tagged removed to rule out a significant role for SUMO in the interaction.

### Matrix Optimization and Score calculation

The Phosphosite O-GlcNAcylation database (1829 peptides) was used as a positive dataset to optimize a scoring matrix. The negative dataset consisted of peptides extracted from IDRs, which we experimentally determined to not be glycosylated by OGT under optimal conditions. It consisted of 135 peptides centered on serines or threonines extracted from these proteins. In-house software was used to optimize the matrix to maximize the scoring difference between the positive and negative datasets, using an iterative process of random changes to the matrix. The final matrix was then used to score glycosylation sites. The threshold for a positive site was set at 160, unless otherwise indicated. (For Table 4, the low, medium and high thresholds are 123, 148 and 160 respectively). All existing predictors suffer from high numbers of false positive predictions. Setting a relatively high threshold increases the likelihood that a positive prediction is accurate, but results in poor sensitivity.

### Measuring Compositional Bias

We measured compositional bias as defined by Harrison and Gerstein(89) and implemented in fLPS2.0(65). The background proportions of amino acid types were those derived from human UniProt records(53) or from a dataset of disordered proteins, determined from the MobiDB(66) manually curated version of the DisProt database(67). We further selected only human proteins with greater than 50% fractional disorder. The PhosphoSite database was used without modification. However, the negative dataset is composed of overlapping peptides and thus highly redundant, which we thought would significantly comprise the bias calculation. Therefore, we used the sequences of the protein regions containing the peptides in the database, rather than the database itself. Only biases related to the whole datasets are reported here.

## Abbreviations

O-GlcNAc: O-linked N-acetylglucosamine
OGT: O-GlcNAc transferase
PSSM: position-specific scoring matrix
PTM: post-translational modification
TPR: tetratricopeptide repeat
FUS: fused in sarcoma
IDR: intrinsically disordered region
LCR_N_: N-terminal low-complexity region

## Data availability

The LC-MS/MS data on EWS-LC and CBP ID3 glycosylation, an optimized scoring matrix and the script for scoring peptides can be downloaded from https://zenodo.org,

DOI: 10.5281/zenodo.6986306.

## Supporting Information

This article contains supporting information.

Table S1. Sequences of WT and mutant FUS LCR_N_.

Figure S1. Amino acid proportions in the human proteome and set of disordered proteins.

Figure S2. ^1^H-^15^N HSQC spectra of WT FUS LCR_N_ and Mut-F FUS LCR_N_ in the presence and absence of OGT-TPR.

## Acknowledgments

We thank Drs. Iva Pritisanac, Tae Hun Kim, Brian Tsang and Robert Vernon for helpful discussions. Oliver Ocsenas and Dr. Rhea Hudson are thanked for help with protein purification. We thank Drs. Suzanne Walker and Peter Tompa for the kind gifts of plasmids for expression of the OGT and CBP intrinsically disordered region proteins respectively. Drs. Paul Taylor and Craig Simpson and The Hospital for Sick Children SPARC Molecular Analysis facility are thanked for their mass spectrometry services. Drs. Cangzhi Jia and Quan Zou are thanked for assistance in providing results of their predictor for select proteins. Funding was provided by grants to J.D.F.-K. from the Canadian Institutes for Health Research (CIHR, FND-148375) and the Natural Sciences and Engineering Research Council of Canada (NSERC, 2016-06718).

## References

1. Torres, C. R., and Hart, G. W. (1984) Topography and polypeptide distribution of terminal N-acetylglucosamine residues on the surfaces of intact lymphocytes. Evidence for O-linked GlcNAc. J. Biol. Chem. 259, 3308–3317

2. Joiner, C. M., Li, H., Jiang, J., and Walker, S. (2019) Structural characterization of the O-GlcNAc cycling enzymes: insights into substrate recognition and catalytic mechanisms. Curr. Opin. Struct. Biol. 56, 97–106

3. Hahne, H., Sobotzki, N., Nyberg, T., Helm, D., Borodkin, V. S., Van Aalten, D. M. F., Agnew, B., and Kuster, B. (2013) Proteome wide purification and identification of O-GlcNAc modified proteins using Click chemistry and mass spectrometry. J. Proteome Res. 12, 927–936

4. Wells, L., Vosseller, K., Cole, R. N., Cronshaw, J. M., Matunis, M. J., and Hart, G. W. (2002) Mapping Sites of O -GlcNAc Modification Using Affinity Tags for Serine and Threonine Post-translational Modifications. Mol. Cell. Proteomics. 1, 791–804

5. Trinidad, J. C., Barkan, D. T., Gulledge, B. F., Thalhammer, A., Sali, A., Schoepfer, R., and Burlingame, A. L. (2012) Global Identification and Characterization of Both O-GlcNAcylation and Phosphorylation at the Murine Synapse. Mol. Cell. Proteomics. 11, 215–229

6. Xu, S.-L., Chalkley, R. J., Maynard, J. C., Wang, W., Ni, W., Jiang, X., Shin, K., Cheng, L., Savage, D., Hühmer, A. F. R., Burlingame, A. L., and Wang, Z.-Y. (2017) Proteomic analysis reveals O-GlcNAc modification on proteins with key regulatory functions in Arabidopsis. Proc. Natl. Acad. Sci. U. S. A. 114, E1536–E1543

7. O’Donnell, N., Zachara, N. E., Hart, G. W., and Marth, J. D. (2004) Ogt-Dependent X-Chromosome-Linked Protein Glycosylation Is a Requisite Modification in Somatic Cell Function and Embryo Viability. Mol. Cell. Biol. 24, 1680–1690

8. Shafi, R., Iyer, S. P. N., Ellies, L. G., O’Donnell, N., Marek, K. W., Chui, D., Hart, G. W., and Marth, J. D. (2000) The O-GlcNAc transferase gene resides on the X chromosome and is essential for embryonic stem cell viability and mouse ontogeny. Proc. Natl. Acad. Sci. U. S. A. 97, 5735–5739

9. Hart, G. W. (2019) Nutrient regulation of signaling and transcription. J. Biol. Chem. 294, 2211–2231

10. Griffith, L. S., Mathes, M., and Schmitz, B. (1995) Beta-amyloid precursor protein is modified with O-linked N-acetylglucosamine. J. Neurosci. Res. 41, 270–8

11. Yuzwa, S. A., Cheung, A. H., Okon, M., Mcintosh, L. P., and Vocadlo, D. J. (2014) O-GlcNAc Modification of tau Directly Inhibits Its Aggregation without Perturbing the Conformational Properties of tau Monomers. J. Mol. Biol. 426, 1736–1752

12. Levine, P. M., Galesic, A., Balana, A. T., Mahul-Mellier, A.-L., Navarro, M. X., De Leon, C. A., Lashuel, H. A., and Pratt, M. R. (2019) α-Synuclein O-GlcNAcylation alters aggregation and toxicity, revealing certain residues as potential inhibitors of Parkinson’s disease. Proc. Natl. Acad. Sci. U. S. A. 116, 1511–1519

13. Sprung, R., Nandi, A., Chen, Y., Kim, S. C., Barma, D., Falck, J. R., and Zhao, Y. (2005) Tagging-via-substrate strategy for probing O-GlcNAc modified proteins. J. Proteome Res. 4, 950–957

14. Pravata, V. M., Gundogdu, M., Bartual, S. G., Ferenbach, A. T., Stavridis, M., Õunap, K., Pajusalu, S., Žordania, R., Wojcik, M. H., and van Aalten, D. M. F. (2020) A missense mutation in the catalytic domain of O-GlcNAc transferase links perturbations in protein O-GlcNAcylation to X-linked intellectual disability. FEBS Lett. 594, 717–727

15. Corfield, A. (2017) Eukaryotic protein glycosylation: a primer for histochemists and cell biologists. Histochem. Cell Biol. 147, 119–147

16. King, D. T., Serrano-Negrón, J. E., Zhu, Y., Moore, C. L., Shoulders, M. D., Foster, L. J., and Vocadlo, D. J. (2022) Thermal Proteome Profiling Reveals the O-GlcNAc-Dependent Meltome. J. Am. Chem. Soc. 144, 3833–3842

17. Yuzwa, S. A., Shan, X., MacAuley, M. S., Clark, T., Skorobogatko, Y., Vosseller, K., and Vocadlo, D. J. (2012) Increasing O-GlcNAc slows neurodegeneration and stabilizes tau against aggregation. Nat. Chem. Biol. 8, 393–399

18. Marotta, N. P., Lin, Y. H., Lewis, Y. E., Ambroso, M. R., Zaro, B. W., Roth, M. T., Arnold, D. B., Langen, R., and Pratt, M. R. (2015) O-GlcNAc modification blocks the aggregation and toxicity of the protein α-synuclein associated with Parkinson’s disease. Nat. Chem. 7, 913–920

19. Nosella, M. L., Tereshchenko, M., Pritišanac, I., Chong, P. A., Toretsky, J. A., Lee, H. O., and Forman-Kay, J. D. (2021) O-Linked-N-Acetylglucosaminylation of the RNA-Binding Protein EWS N-Terminal Low Complexity Region Reduces Phase Separation and Enhances Condensate Dynamics. J. Am. Chem. Soc. 143, 11520–11534

20. Kim, T. H., Payliss, B. J., Nosella, M. L., Lee, I. T. W., Toyama, Y., Forman-Kay, J. D., and Kay, L. E. (2021) Interaction hot spots for phase separation revealed by NMR studies of a CAPRIN1 condensed phase. Proc. Natl. Acad. Sci. U. S. A. 118, e2104897118

21. Lv, P., Du, Y., He, C., Peng, L., Zhou, X., Wan, Y., Zeng, M., Zhou, W., Zou, P., Li, C., Zhang, M., Dong, S., and Chen, X. (2022) O-GlcNAcylation modulates liquid-liquid phase separation of SynGAP/PSD-95. Nat. Chem. 10.1038/s41557-022-00946-9

22. Pathak, S., Alonso, J., Schimpl, M., Rafie, K., Blair, D. E., Borodkin, V. S., Schüttelkopf, A. W., Albarbarawi, O., and van Aalten, D. M. F. (2015) The active site of O-GlcNAc transferase imposes constraints on substrate sequence. Nat. Struct. Mol. Biol. 22, 744–750

23. Wu, H. Y., Lu, C. T., Kao, H. J., Chen, Y. J., Chen, Y. J., and Lee, T. Y. (2014) Characterization and identification of protein O-GlcNAcylation sites with substrate specificity. BMC Bioinformatics. 15, 1–12

24. Jinek, M., Rehwinkel, J., Lazarus, B. D., Izaurralde, E., Hanover, J. A., and Conti, E. (2004) The superhelical TPR-repeat domain of O-linked GlcNAc transferase exhibits structural similarities to importin alpha. Nat Struct Mol Biol. 11, 1001–1007

25. Leney, A. C., El Atmioui, D., Wu, W., Ovaa, H., and Heck, A. J. R. (2017) Elucidating crosstalk mechanisms between phosphorylation and O-GlcNAcylation. Proc. Natl. Acad. Sci. U. S. A. 114, E7255–E7261

26. Ramirez, D. H., Yang, B., D’Souza, A. K., Shen, D., and Woo, C. M. (2021) Truncation of the TPR domain of OGT alters substrate and glycosite selection. Anal. Bioanal. Chem. 413, 7385–7399

27. Ortiz-Meoz, R. F., Merbl, Y., Kirschner, M. W., and Walker, S. (2014) Microarray discovery of new OGT substrates: The medulloblastoma oncogene OTX2 is O -GlcNAcylated. J. Am. Chem. Soc. 136, 4845–4848

28. Gao, Y., Wells, L., Comer, F. I., Parker, G. J., and Hart, G. W. (2001) Dynamic O-glycosylation of nuclear and cytosolic proteins: cloning and characterization of a neutral, cytosolic beta-N-acetylglucosaminidase from human brain. J. Biol. Chem. 276, 9838–45

29. Hanover, J. A., Yu, S., Lubas, W. B., Shin, S. H., Ragano-Caracciola, M., Kochran, J., and Love, D. C. (2003) Mitochondrial and nucleocytoplasmic isoforms of O-linked GlcNAc transferase encoded by a single mammalian gene. Arch. Biochem. Biophys. 409, 287–97

30. Lu, L., Fan, D., Hu, C. W., Worth, M., Ma, Z. X., and Jiang, J. (2016) Distributive O-GlcNAcylation on the Highly Repetitive C-Terminal Domain of RNA Polymerase II. Biochemistry. 55, 1149–1158

31. Iyer, S. P. N., and Hart, G. W. (2003) Roles of the Tetratricopeptide Repeat Domain in O-GlcNAc Transferase Targeting and Protein Substrate Specificity. J. Biol. Chem. 278, 24608–24616

32. Lubas, W. A., and Hanover, J. A. (2000) Functional Expression of O -linked GlcNAc Transferase. J. Biol. Chem. 275, 10983–10988

33. Yang, X., Zhang, F., and Kudlow, J. E. (2002) Recruitment of O-GlcNAc transferase to promoters by corepressor mSin3A: Coupling protein O-GlcNAcylation to transcriptional repression. Cell. 110, 69–80

34. Zhang, Q., Liu, X., Gao, W., Li, P., Hou, J., Li, J., and Wong, J. (2014) Differential regulation of the ten-eleven translocation (TET) family of dioxygenases by O-linked β-N-acetylglucosamine transferase (OGT). J. Biol. Chem. 289, 5986–5996

35. Cheung, W. D., Sakabe, K., Housley, M. P., Dias, W. B., and Hart, G. W. (2008) O-linked β-N-acetylglucosaminyltransferase substrate specificity is regulated by myosin phosphatase targeting and other interacting proteins. J. Biol. Chem. 283, 33935–33941

36. Wani, W. Y., Chatham, J. C., Darley-Usmar, V., McMahon, L. L., and Zhang, J. (2017) O-GlcNAcylation and neurodegeneration. Brain Res. Bull. 133, 80–87

37. Hart, G. W., Housley, M. P., and Slawson, C. (2007) Cycling of O-linked β-N-acetylglucosamine on nucleocytoplasmic proteins. Nature. 446, 1017–1022

38. Hornbeck, P. V, Kornhauser, J. M., Tkachev, S., Zhang, B., Skrzypek, E., Murray, B., Latham, V., and Sullivan, M. (2012) PhosphoSitePlus: a comprehensive resource for investigating the structure and function of experimentally determined post-translational modifications in man and mouse. Nucleic Acids Res. 40, D261–70

39. Li, F., Li, C., Revote, J., Zhang, Y., Webb, G. I., Li, J., Song, J., and Lithgow, T. (2016) GlycoMinestruct: A new bioinformatics tool for highly accurate mapping of the human N-linked and O-linked glycoproteomes by incorporating structural features. Sci. Rep. 6, 1–16

40. Li, F., Li, C., Wang, M., Webb, G. I., Zhang, Y., Whisstock, J. C., and Song, J. (2015) GlycoMine: A machine learning-based approach for predicting N-, C-and O-linked glycosylation in the human proteome. Bioinformatics. 31, 1411–1419

41. Jia, C., Zuo, Y., and Zou, Q. (2018) O-GlcNAcPRED-II: An integrated classification algorithm for identifying O-GlcNAcylation sites based on fuzzy undersampling and a K -means PCA oversampling technique. Bioinformatics. 34, 2029–2036

42. Kao, H. J., Huang, C. H., Bretaña, N. A., Lu, C. T., Huang, K. Y., Weng, S. L., and Lee, T. Y. (2015) A two-layered machine learning method to identify protein O-GlcNAcylation sites with O-GlcNAc transferase substrate motifs. BMC Bioinformatics. 16, 1–11

43. Gupta, R., and Brunak, S. (2002) Prediction of glycosylation across the human proteome and the correlation to protein function. Pac. Symp. Biocomput. 322, 310–22

44. Wang, J., Torii, M., Liu, H., Hart, G. W., and Hu, Z.-Z. (2011) dbOGAP - an integrated bioinformatics resource for protein O-GlcNAcylation. BMC Bioinformatics. 12, 91

45. Mauri, T., Menu-Bouaouiche, L., Bardor, M., Lefebvre, T., Lensink, M. F., and Brysbaert, G. (2021) O-GlcNAcylation Prediction: An Unattained Objective. Adv. Appl. Bioinform. Chem. 14, 87–102

46. Chauhan, J. S., Rao, A., and Raghava, G. P. S. (2013) In silico Platform for Prediction of N-, O-and C-Glycosites in Eukaryotic Protein Sequences. PLoS One. 8, 1–10

47. Simon, D. N., Wriston, A., Fan, Q., Shabanowitz, J., Florwick, A., Dharmaraj, T., Peterson, S. B., Gruenbaum, Y., Carlson, C. R., Grønning-Wang, L. M., Hunt, D. F., and Wilson, K. L. (2018) OGT (O-GlcNAc Transferase) Selectively Modifies Multiple Residues Unique to Lamin A. Cells. 7, 44

48. Jochmann, R., Holz, P., Sticht, H., and Stürzl, M. (2014) Validation of the reliability of computational O-GlcNAc prediction. Biochim. Biophys. Acta - Proteins Proteomics. 1844, 416–421

49. Lazarus, M. B., Nam, Y., Jiang, J., Sliz, P., and Walker, S. (2011) Structure of human O-GlcNAc transferase and its complex with a peptide substrate. Nature. 469, 564–569

50. Schimpl, M., Zheng, X., Borodkin, V. S., Blair, D. E., Ferenbach, A. T., Schüttelkopf, A. W., Navratilova, I., Aristotelous, T., Albarbarawi, O., Robinson, D. A., MacNaughtan, M. A., and Van Aalten, D. M. F. (2012) O-GlcNAc transferase invokes nucleotide sugar pyrophosphate participation in catalysis. Nat. Chem. Biol. 8, 969–974

51. Joiner, C. M., Levine, Z. G., Aonbangkhen, C., Woo, C. M., and Walker, S. (2019) Aspartate residues far from the active site drive O-GlcNAc transferase substrate selection. J. Am. Chem. Soc. 141, 12974–12978

52. Levine, Z. G., Fan, C., Melicher, M. S., Orman, M., Benjamin, T., and Walker, S. (2018) O -GlcNAc Transferase Recognizes Protein Substrates Using an Asparagine Ladder in the Tetratricopeptide Repeat (TPR) Superhelix. J. Am. Chem. Soc. 140, 3510–3513

53. UniProt Consortium (2021) UniProt: the universal protein knowledgebase in 2021. Nucleic Acids Res. 49, D480–D489

54. Ramirez, D. H., Aonbangkhen, C., Wu, H., Naftaly, J. A., Tang, S., O’Meara, T. R., and Woo, C. M. (2020) Engineering a Proximity-Directed O-GlcNAc Transferase for Selective Protein O-GlcNAcylation in Cells. ACS Chem. Biol. 15, 1059–1066

55. Bantscheff, M., Lemeer, S., Savitski, M. M., and Kuster, B. (2012) Quantitative mass spectrometry in proteomics: Critical review update from 2007 to the present. Anal. Bioanal. Chem. 404, 939–965

56. Greis, K., and Hart, G. W. (1998) Analytical methods for the study of O-GlcNAc glycoproteins and glycopeptides. Methods Mol. Biol. 76, 19–33

57. Wang, Z., Udeshi, N. D., O’Malley, M., Shabanowitz, J., Hunt, D. F., and Hart, G. W. (2010) Enrichment and site mapping of O-linked N-acetylglucosamine by a combination of chemical/enzymatic tagging, photochemical cleavage, and electron transfer dissociation mass spectrometry. Mol. Cell. Proteomics. 9, 153–160

58. Kamemura, K. (2016) O-GlcNAc glycosylation stoichiometry of the FET protein family: only EWS is glycosylated with a high stoichiometry. Biosci. Biotechnol. Biochem. 8451, 1–6

59. Bachmaier, R., Aryee, D. N. T., Jug, G., Kauer, M., Kreppel, M., Lee, K. A. W., and Kovar, H. (2009) O-GlcNAcylation is involved in the transcriptional activity of EWS-FLI1 in Ewing’s sarcoma. Oncogene. 28, 1280–1284

60. Chong, P. A., Ozdamar, B., Wrana, J. L., and Forman-Kay, J. D. (2004) Disorder in a target for the smad2 mad homology 2 domain and its implications for binding and specificity. J. Biol. Chem. 279, 40707–14

61. Nott, T. J., Petsalaki, E., Farber, P., Jervis, D., Fussner, E., Plochowietz, A., Craggs, T. D., Bazett-Jones, D. P., Pawson, T., Forman-Kay, J. D., and Baldwin, A. J. (2015) Phase transition of a disordered nuage protein generates environmentally responsive membraneless organelles. Mol. Cell. 57, 936–947

62. Baker, J. M. R., Hudson, R. P., Kanelis, V., Choy, W.-Y., Thibodeau, P. H., Thomas, P. J., and Forman-Kay, J. D. (2007) CFTR regulatory region interacts with NBD1 predominantly via multiple transient helices. Nat. Struct. Mol. Biol. 14, 738–45

63. Tsang, B., Arsenault, J., Vernon, R. M., Lin, H., Sonenberg, N., Wang, L., Bah, A., and Forman-Kay, J. D. (2019) Phosphoregulated FMRP phase separation models activity-dependent translation through bidirectional control of mRNA granule formation. Proc. Natl. Acad. Sci. U. S. A. 116, 4218–4227

64. Mittag, T., Orlicky, S., Choy, W. Y., Tang, X., Lin, H., Sicheri, F., Kay, L. E., Tyers, M., and Forman-Kay, J. D. (2008) Dynamic equilibrium engagement of a polyvalent ligand with a single-site receptor. Proc Natl Acad Sci U S A. 105, 17772–17777

65. Harrison, P. M. (2021) fLPS 2.0: Rapid annotation of compositionally-biased regions in biological sequences. PeerJ. 10.7717/peerj.12363

66. Piovesan, D., Necci, M., Escobedo, N., Monzon, A. M., Hatos, A., Mičetić, I., Quaglia, F., Paladin, L., Ramasamy, P., Dosztányi, Z., Vranken, W. F., Davey, N. E., Parisi, G., Fuxreiter, M., and Tosatto, S. C. E. (2021) MobiDB: intrinsically disordered proteins in 2021. Nucleic Acids Res. 49, D361–D367

67. Quaglia, F., Mészáros, B., Salladini, E., Hatos, A., Pancsa, R., Chemes, L. B., Pajkos, M., Lazar, T., Peña-Díaz, S., Santos, J., Ács, V., Farahi, N., Fichó, E., Aspromonte, M. C., Bassot, C., Chasapi, A., Davey, N. E., Davidović, R., Dobson, L., Elofsson, A., Erdős, G., Gaudet, P., Giglio, M., Glavina, J., Iserte, J., Iglesias, V., Kálmán, Z., Lambrughi, M., Leonardi, E., Longhi, S., Macedo-Ribeiro, S., Maiani, E., Marchetti, J., Marino-Buslje, C., Mészáros, A., Monzon, A. M., Minervini, G., Nadendla, S., Nilsson, J. F., Novotný, M., Ouzounis, C. A., Palopoli, N., Papaleo, E., Pereira, P. J. B., Pozzati, G., Promponas, V. J., Pujols, J., Rocha, A. C. S., Salas, M., Sawicki, L. R., Schad, E., Shenoy, A., Szaniszló, T., Tsirigos, K. D., Veljkovic, N., Parisi, G., Ventura, S., Dosztányi, Z., Tompa, P., Tosatto, S. C. E., and Piovesan, D. (2022) DisProt in 2022: improved quality and accessibility of protein intrinsic disorder annotation. Nucleic Acids Res. 50, D480–D487

68. Bozoky, Z., Krzeminski, M., Muhandiram, R., Birtley, J. R., Al-Zahrani, A., Thomas, P. J., Frizzell, R. A., Ford, R. C., and Forman-Kay, J. D. (2013) Regulatory R region of the CFTR chloride channel is a dynamic integrator of phospho-dependent intra-and intermolecular interactions. Proc. Natl. Acad. Sci. U. S. A. 110, E4427–36

69. Contreras-Martos, S., Piai, A., Kosol, S., Varadi, M., Bekesi, A., Lebrun, P., Volkov, A. N., Gevaert, K., Pierattelli, R., Felli, I. C., and Tompa, P. (2017) Linking functions: An additional role for an intrinsically disordered linker domain in the transcriptional coactivator CBP. Sci. Rep. 7, 1–13

70. Piai, A., Calçada, E. O., Tarenzi, T., Grande, A. Del, Varadi, M., Tompa, P., Felli, I. C., and Pierattelli, R. (2016) Just a Flexible Linker? the Structural and Dynamic Properties of CBP-ID4 Revealed by NMR Spectroscopy. Biophys. J. 110, 372–381

71. Kosol, S., Contreras-Martos, S., Piai, A., Varadi, M., Lazar, T., Bekesi, A., Lebrun, P., Felli, I. C., Pierattelli, R., and Tompa, P. (2020) Interaction between the scaffold proteins CBP by IQGAP1 provides an interface between gene expression and cytoskeletal activity. Sci. Rep. 10, 5753

72. Lazarus, M. B., Jiang, J., Kapuria, V., Bhuiyan, T., Janetzko, J., Zandberg, W. F., Vocadlo, D. J., Herr, W., and Walker, S. (2013) HCF-1 is cleaved in the active site of O-GlcNAc transferase. Science. 342, 1235–9

73. Kapuria, V., Röhrig, U. F., Waridel, P., Lammers, F., Borodkin, V. S., Van Aalten, D. M. F., Zoete, V., and Herr, W. (2018) The conserved threonine-rich region of the HCF-1PRO repeat activates promiscuous OGT:UDP-GlcNAc glycosylation and proteolysis activities. J. Biol. Chem. 293, 17754–17768

74. Balana, A. T., Mukherjee, A., Nagpal, H., Moon, S. P., Fierz, B., Vasquez, K. M., and Pratt, M. R. (2021) O-GlcNAcylation of High Mobility Group Box 1 (HMGB1) Alters Its DNA Binding and DNA Damage Processing Activities. J. Am. Chem. Soc. 143, 16030–16040

75. Fujiki, R., Hashiba, W., Sekine, H., Yokoyama, A., Chikanishi, T., Ito, S., Imai, Y., Kim, J., He, H. H., Igarashi, K., Kanno, J., Ohtake, F., Kitagawa, H., Roeder, R. G., Brown, M., and Kato, S. (2011) GlcNAcylation of histone H2B facilitates its monoubiquitination. Nature. 480, 557–560

76. Roquemore, E. P., Chevrier, M. R., Cotter, R. J., and Hart, G. W. (1996) Dynamic O-GlcNAcylation of the small heat shock protein αB-crystallin. Biochemistry. 35, 3578–3586

77. Meek, R. W., Blaza, J. N., Busmann, J. A., Alteen, M. G., Vocadlo, D. J., and Davies, G. J. (2021) Cryo-EM structure provides insights into the dimer arrangement of the O-linked β-N-acetylglucosamine transferase OGT. Nat. Commun. 10.1038/s41467-021-26796-6

78. Springhower, C. E., Rosen, M. K., and Chook, Y. M. (2020) Karyopherins and condensates. Curr. Opin. Cell Biol. 64, 112–123

79. Zhu, Y., Willems, L. I., Salas, D., Cecioni, S., Wu, W. B., Foster, L. J., and Vocadlo, D. J. (2020) Tandem Bioorthogonal Labeling Uncovers Endogenous Cotranslationally O-GlcNAc Modified Nascent Proteins. J. Am. Chem. Soc. 142, 15729–15739

80. Zhu, Y., Liu, T. W., Cecioni, S., Eskandari, R., Zandberg, W. F., and Vocadlo, D. J. (2015) O-GlcNAc occurs cotranslationally to stabilize nascent polypeptide chains. Nat. Chem. Biol. 11, 319–325

81. Jumper, J., Evans, R., Pritzel, A., Green, T., Figurnov, M., Ronneberger, O., Tunyasuvunakool, K., Bates, R., Žídek, A., Potapenko, A., Bridgland, A., Meyer, C., Kohl, S. A. A., Ballard, A. J., Cowie, A., Romera-Paredes, B., Nikolov, S., Jain, R., Adler, J., Back, T., Petersen, S., Reiman, D., Clancy, E., Zielinski, M., Steinegger, M., Pacholska, M., Berghammer, T., Bodenstein, S., Silver, D., Vinyals, O., Senior, A. W., Kavukcuoglu, K., Kohli, P., and Hassabis, D. (2021) Highly accurate protein structure prediction with AlphaFold. Nature. 596, 583–589

82. Baek, M., DiMaio, F., Anishchenko, I., Dauparas, J., Ovchinnikov, S., Lee, G. R., Wang, J., Cong, Q., Kinch, L. N., Schaeffer, R. D., Millán, C., Park, H., Adams, C., Glassman, C. R., DeGiovanni, A., Pereira, J. H., Rodrigues, A. V, van Dijk, A. A., Ebrecht, A. C., Opperman, D. J., Sagmeister, T., Buhlheller, C., Pavkov-Keller, T., Rathinaswamy, M. K., Dalwadi, U., Yip, C. K., Burke, J. E., Garcia, K. C., Grishin, N. V, Adams, P. D., Read, R. J., and Baker, D. (2021) Accurate prediction of protein structures and interactions using a three-track neural network. Science. 373, 871–876

83. Kay, L. E., Keifer, P., and Saarinen, T. (1992) Pure Absorption Gradient Enhanced Heteronuclear Single Quantum Correlation Spectroscopy with Improved Sensitivity. J. Am. Chem. Soc. 114, 10663–10665

84. Delaglio, F., Grzesiek, S., Vuister, G. W., Zhu, G., Pfeifer, J., and Bax, A. (1995) NMRPipe: a multidimensional spectral processing system based on UNIX pipes. J. Biomol. NMR. 6, 277–293

85. Vranken, W. F., Boucher, W., Stevens, T. J., Fogh, R. H., Pajon, A., Llinas, M., Ulrich, E. L., Markley, J. L., Ionides, J., and Laue, E. D. (2005) The CCPN data model for NMR spectroscopy: development of a software pipeline. Proteins. 59, 687–696

86. Lee, W., Tonelli, M., and Markley, J. L. (2015) NMRFAM-SPARKY: enhanced software for biomolecular NMR spectroscopy. Bioinformatics. 31, 1325–1327

87. Ulrich, E. L., Akutsu, H., Doreleijers, J. F., Harano, Y., Ioannidis, Y. E., Lin, J., Livny, M., Mading, S., Maziuk, D., Miller, Z., Nakatani, E., Schulte, C. F., Tolmie, D. E., Kent Wenger, R., Yao, H., and Markley, J. L. (2008) BioMagResBank. Nucleic Acids Res. 36, D402–8

88. Burke, K. A., Janke, A. M., Rhine, C. L., and Fawzi, N. L. (2015) Residue-by-Residue View of In Vitro FUS Granules that Bind the C-Terminal Domain of RNA Polymerase II. Mol. Cell. 60, 1–11

89. Harrison, P. M., and Gerstein, M. (2003) A method to assess compositional bias in biological sequences and its application to prion-like glutamine/asparagine-rich domains in eukaryotic proteomes. Genome Biol.

