## Supplemental file for "Exploration of O-GlcNAc-transferase (OGT) glycosylation sites reveals a target sequence compositional bias"

Table S1. Sequences of WT and mutant FUS $LCR_N$. Mutated positions are highlighted in red.

| Protein | Sequence |
|---|---|
| WT | MASNDYTQQA TQSYGAYPTQ PGQGYSQQSS QPYGQQSYSG YSQSTDTSGY GQSSYSSYGQ SQNTGYGTQS TPQGYGSTGG YGSSQSSQSS YGQQSSYPGY GQQPAPSSTS GSYGSSSQSS SYGQPQSGSY SQQPSYGGQQ QSYGQQQSYN PPQGYGQQNQ YNSSSGGGGG GGGGGNYGQD QSSMSSGGGS GGGYGNQDQS GGGGSGGYGQ QDRG |
| Mut-A | MASNDYTQQA TQSYGAYPTQ PGQGYSQQSS **T**PY**A**Q**A**SYS**T** YS**P**ST**T**TS**A**Y GQSSYSSYGQ SQNTGYGTQS TPQGYGSTGG YGSSQSSQSS YGQQSSYPGY GQQPAPSSTS GSYGSSSQSS SYGQPQSGSY SQQPSYGGQQ QSYGQQQSYN PPQGYGQQNQ YNSSSGGGGG GGGGGNYGQD QSSMSSGGGS GGGYGNQDQS GGGGSGGYGQ QDRG |
| Mut-B | MASNDYTQQA TQSYGAYPTQ PGQGYSQQSS QPYGQQSYSG YSQSTDTSGY GQSSYSSYGQ SQNTGY**A**T**T**S TPQ**A**Y**P**ST**AP** Y**AT**S**P**SSQSS YGQQSSYPGY GQQPAPSSTS GSYGSSSQSS SYGQPQSGSY SQQPSYGGQQ QSYGQQQSYN PPQGYGQQNQ YNSSSGGGGG GGGGGNYGQD QSSMSSGGGS GGGYGNQDQS GGGGSGGYGQ QDRG |
| Mut-C | MASNDYTQQA TQSYGAYPTQ PGQGYSQQSS QPYGQQSYSG YSQSTDTSGY GQSSYSSYGQ SQNTGYGTQS TPQGYGSTGG YGSSQSSQSS YGQQSSYP**A**Y **T**QQPAP**TT**TS **A**SY**AT**SSQSS SYGQPQSGSY SQQPSYGGQQ QSYGQQQSYN PPQGYGQQNQ YNSSSGGGGG GGGGGNYGQD QSSMSSGGGS GGGYGNQDQS GGGGSGGYGQ QDRG |
| Mut-D | MASNDYTQQA TQSYGAYPTQ PGQGYSQQSS **T**PY**A**Q**A**SYS**T** YS**P**ST**T**TS**A**Y GQSSYSSYGQ SQNTGY**A**T**T**S TPQ**A**Y**P**ST**AP** Y**AT**S**P**SSQSS YGQQSSYPGY GQQPAPSSTS GSYGSSSQSS SYGQPQSGSY SQQPSYGGQQ QSYGQQQSYN PPQGYGQQNQ YNSSSGGGGG GGGGGNYGQD QSSMSSGGGS GGGYGNQDQS GGGGSGGYGQ QDRG |
| Mut-E | MASNDYTQQA TQSYGAYPTQ PGQGYSQQSS QPYGQQSYSG YSQSTDTSGY GQSSYSSYGQ SQNTGY**A**T**T**S TPQ**A**Y**P**ST**AP** Y**AT**S**P**SSQSS YGQQSSYP**A**Y **T**QQPAP**TT**TS **A**SY**AT**SSQSS SYGQPQSGSY SQQPSYGGQQ QSYGQQQSYN PPQGYGQQNQ YNSSSGGGGG GGGGGNYGQD QSSMSSGGGS GGGYGNQDQS GGGGSGGYGQ QDRG |
| Mut-F | MASNDYTQQA TQSYGAYPTQ PGQGYSQQSS **T**PY**A**Q**A**SYS**T** YS**P**ST**T**TS**A**Y GQSSYSSYGQ SQNTGY**A**T**T**S TPQ**A**Y**P**ST**AP** Y**AT**S**P**SSQSS YGQQSSYP**A**Y **T**QQPAP**TT**TS **A**SY**AT**SSQSS SYGQPQSGSY SQQPSYGGQQ QSYGQQQSYN PPQGYGQQNQ YNSSSGGGGG GGGGGNYGQD QSSMSSGGGS GGGYGNQDQS GGGGSGGYGQ QDRG |

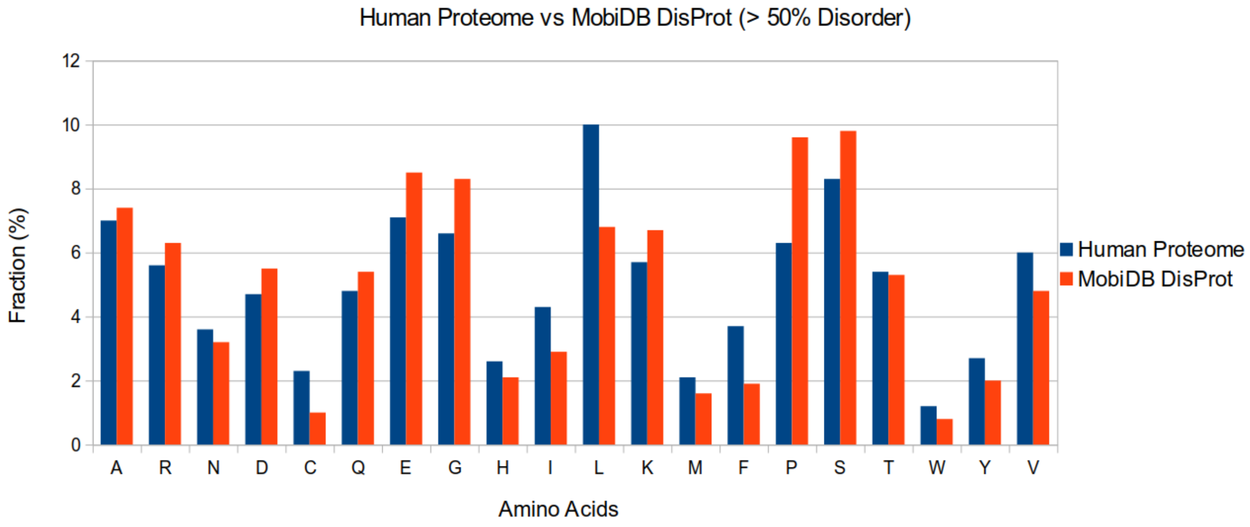


Figure S1: Amino acid proportions in the human proteome and set of disordered proteins. For the human proteome, we used all human proteins in the Uniprot database. For the disordered set, we used a manually curated MobiDB version of the DisProt database and selected for human proteins with greater than 50% defined fractional disorder content.

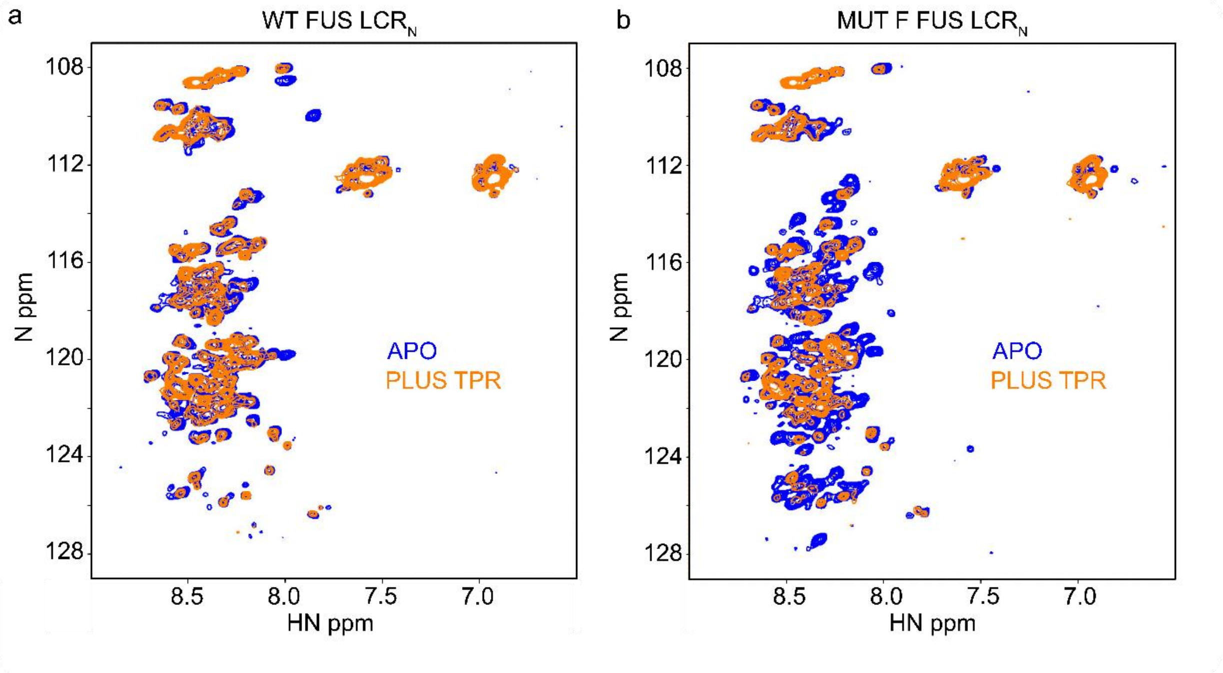


Figure S2: $^1H$-$^{15}N$ HSQC spectra of WT FUS LCR$_N$ (a) and Mut-F FUS LCR$_N$ (b) in the presence and absence of OGT-TPR. Spectra of the FUS LCR$_N$ at 20 μM are shown in blue and spectra in the presence of 50 μM OGT-TPR are shown in orange. Spectra were recorded with a field strength of 600 MHz at 5°C in a buffer comprised of 40 mM $KPO_4$, 125 mM NaCl, 0.5 mM EDTA, 0.5 mM benzamidine, 5 mM DTT and 10% $D_2O$, pH 7.2.
